# Intestinal helminth infection impairs oral and parenteral vaccine efficacy

**DOI:** 10.1101/2022.09.22.508369

**Authors:** LaKeya C. Hardy, Camille M. Kapita, Evelyn Campbell, Jason A. Hall, Joseph F. Urban, Yasmine Belkaid, Cathryn R. Nagler, Onyinye I. Iweala

## Abstract

The impact of endemic parasitic infection on vaccine efficacy is an important consideration for vaccine development and deployment. We have examined whether intestinal infection with the natural murine helminth *Heligmosomoides polygyrus bakeri* alters antigen-specific antibody and cellular immune responses to oral and parenteral vaccination in mice. We found that oral vaccination of mice with a clinically relevant, live, attenuated, recombinant *Salmonella* vaccine that expresses chicken egg ovalbumin (*Salmonella*-OVA) disrupts ovalbumin-specific regulatory T cell networks in the gut associated lymphoid tissue and promotes T-effector responses to OVA. Chronic intestinal helminth infection significantly reduced Th1-skewed antibody responses to oral vaccination with *Salmonella-*OVA. Activated, adoptively-transferred, OVA-specific CD4^+^ T cells accumulated in draining mesenteric lymph nodes (MLN) of vaccinated mice, irrespective of their helminth-infection status. However, helminth infection increased the frequencies of adoptively-transferred OVA-specific CD4^+^ T cells producing IL-4 and IL-10 in the MLN. Chronic intestinal helminth infection also significantly reduced Th2-skewed antibody responses to parenteral vaccination with OVA adsorbed to alum. These findings suggest helminth-induced impairment of vaccine antibody responses may be driven by the development of IL-10-secreting CD4^+^ T regulatory cells. They also underscore the potential need to treat parasitic infection before mass vaccination campaigns in helminth-endemic areas.

## INTRODUCTION

Vaccination is one of the most effective public health measures against infection (1–4). However, as the COVID-19 pandemic has highlighted, there are significant unmet needs for vaccine coverage for adults and children across the world, particularly in low- and middle-income countries where populations shoulder a significant portion of the world’s infectious disease burden (3–5). The World Health Organization (WHO) and other multinational, public-private partnership organizations continue to advocate for strategies that address the availability, affordability, storage and handling, ease of administration, and safety of vaccines, with the goal of expanding vaccination coverage amongst underserved populations (4, 6, 7).

Mucosal vaccines, including oral vaccines, are potent inducers of local mucosal and systemic cellular and humoral immune responses (8–10). Recombinant oral *Salmonella* vaccines, for example, are used in veterinary medicine, especially in the context of poultry farming, to improve fowl health and ensure food safety (11, 12). Oral vaccination with live-attenuated *Salmonella* strains is also used to protect humans against typhoid (13, 14) and paratyphoid fever (15). In humans, oral *Salmonella* vaccines activate circulating B and T cells, expand the number of circulating CD4^+^ Th1 cells, increase serum IFN-γ and TNF-α, and induce *Salmonella*-specific serum and fecal antibody responses (14). In mice, T and B cells are also critical for protective immune responses to attenuated and virulent *Salmonella* (16). CD4^+^ Th1 cells and robust IFN-γ production (17, 18) coupled with *Salmonella*-specific IgG and IgA responses are critical for the clearance of *Salmonella* and development of protective immunity against virulent *Salmonella* strains (18, 19).

One challenge to vaccination is the impact that parasitic gastrointestinal helminth infections can have on the immune response to vaccines (20–22). Greater than 50% of the world’s population lives in regions where helminth infections are endemic (23). Nearly 1.5 billion people are chronically infected with gastrointestinal helminths (23, 24). Three hundred million of these individuals also suffer from malnutrition, stunted growth, anemia, and reduced protective immunity to unrelated pathogens (25, 26). We and others have shown that preexisting helminth infection is a potent modulator of the immune response to orally delivered dietary antigens (27–29), gastrointestinal bacterial infection (30) and parenteral vaccination (22, 31, 32). Epidemiological studies, clinical trials, and animal models suggest that helminth infection has powerful immunosuppressive effects on the development of allergic, autoimmune, and inflammatory diseases (33–35). Helminth infection promotes immune suppression by inducing regulatory cells and cytokines that modulate Th1-, Th2-, and Th17-dependent immune responses (36, 37) in part through interactions with endogenous microbiota (38, 39). Since the regions where helminth infection is endemic overlap significantly with the regions of the world targeted by global health organizations for improved vaccine coverage (1, 4, 40), the impact of helminth infections on vaccine-induced protective immunity must be considered in vaccine design and deployment.

We used a live attenuated oral *Salmonella* vaccine strain expressing chicken egg ovalbumin (OVA) (41) to examine the impact of chronic intestinal helminth infection with the natural mouse parasite *Heligmosomoides polygyrus bakeri* (*H. polygyrus bakeri*) on vaccine antigen-specific cellular and antibody responses. We found that a live attenuated oral *Salmonella*-OVA vaccine disrupts OVA-specific regulatory T cell expansion, promoting OVA-specific T-effector responses. Chronic intestinal helminth infection significantly reduced Th1-skewed antibody responses to oral vaccination with *Salmonella-*OVA even though activated OVA-specific CD4^+^ T cells accumulated in draining mesenteric lymph nodes (MLNs) of helminth-free and helminth-infected mice. Helminth infection also increased the frequencies of adoptively-transferred, OVA-specific CD4^+^ T cells producing IL-4 and IL-10 in the draining MLN. This suggests that IL-10-secreting CD4^+^ T regulatory cells may reduce vaccine-induced antibody responses in helminth-infected mice and highlights the potential need to eliminate immunosuppressive intestinal parasites prior to vaccination in regions where helminth infection is endemic.

## MATERIALS AND METHODS

### Mice

To evaluate peripheral conversion of OVA-specific CD4^+^ T cells to Foxp3^+^ Treg cells, Ly5.1^+^ RAG-1 replete B6.SJL mice and C57BL/6 OT-II transgenic (Tg) RAG-1 KO Ly5.2^+^ mice were purchased from Taconic Farms. Foxp3 eGFP reporter mice (Foxp3^eGFP^) were originally obtained from M. Oukka (Brigham and Women’s Hospital, Cambridge, MA (42)). OT-II Tg RAG-1 KO Ly5.2^+^ Foxp3^eGFP^ mice were generated by crossing the F1 progeny of C57BL/6 OT-II Tg RAG-1 KO Ly5.2^+^ x Foxp3 ^eGFP^ breeders. These mice were maintained at an American Association for the Accreditation of Laboratory Animal Care–accredited animal facility at the National Institute for Allergy and Infectious Diseases (NIAID) and housed following procedures outlined in the Guide for the Care and Use of Laboratory Animals under an animal study proposal approved by the NIAID Animal Care and Use Committee.

For the helminth infection and oral and intramuscular vaccination experiments conducted at Massachusetts General Hospital (MGH), six-to eight-week-old male and female C57BL/6 J mice were purchased from the Jackson Laboratory (Bar Harbor, ME). OT-II (Thy1.1) mice on the C57BL/6 background, transgenic for the TCR recognizing OVA peptide 323-339 were provided by A. Luster (Massachusetts General Hospital (MGH), Charlestown, MA). Mice were fed autoclaved food and water and maintained in a specific-pathogen-free facility at MGH. All experiments were conducted after approval and according to regulations of the Subcommittee on Research Animal Care at MGH.

For helminth infection and intramuscular vaccination experiments conducted at University of North Carolina at Chapel Hill (UNC), eight to sixteen-week-old male and female C57BL/6 J mice were also purchased from the Jackson Laboratory (Bar Harbor, ME). Mice were fed autoclaved food and water and maintained in a specific-pathogen-free facility at the UNC. All mouse experimental procedures were approved by the UNC Institutional Animal Care and Use Committee.

### *In vivo* cell transfer and dietary oral antigen administration

T lymphocytes were extracted from the peripheral LNs (excluding the spleen) of OT-II Tg RAG-1 KO Foxp3^eGFP^ mice (Ly5.2^+^) and adoptively transferred into B6.SJL recipient mice (Ly5.1^+^). Each mouse received 10^6^ cells. Recipient mice were split into two groups. Select groups received a 1.5% OVA solution in drinking water replaced every 48 h (grade V; Sigma-Aldrich) for five consecutive days. The other groups received normal drinking water. On day 6, mesenteric lymph nodes (MLNs – pooled portal, duodenum, jejunum, and ileum LNs as previously described (43)) and intestinal lamina propria (LP) were collected from B6.SJL hosts, and Foxp3-eGFP expression assessed in transferred cells. LN and LP single-cell suspensions were prepared as previously described (44).

### *S. typhimurium* vaccine strains and oral immunization

The recombinant attenuated vaccine strain *Salmonella typhimurium* SL3261 (*aroA,* (45)) carrying either the plasmid pnirOVA (*Salmonella*-OVA) or pnirBEM (*Salmonella-*BEM) were grown overnight shaking at 37°C in Luria Bertani (LB) broth supplemented with 100μg/ml of ampicillin (Ap) as previously described (41). An OD_600_ of 0.5 was estimated to have 2 x 10^8^ bacteria per ml of culture. In experiments examining the peripheral conversion of OVA-specific CD4^+^ T cells to Foxp3^+^ Tregs, subsets of mice were gavaged with 10^12^ attenuated *Salmonella* in PBS one day after congenic cell adoptive transfer. In the helminth-infection experiments, subsets of *H. polygyrus bakeri-*infected and uninfected mice were given 2 to 6 x 10^10^ attenuated *Salmonella* in PBS intragastrically using a 20-gauge ball-tipped feeding needle at different time points (14 and 21 days after parasite inoculation for antibody production experiments, and 7 days after parasite inoculation for cellular immune response experiments). To determine CFU *Salmonella* per gram tissue, spleens were weighed, homogenized in Hanks Balanced Salt Solution and plated on LB plates containing 100 μg/ml ampicillin.

### Intramuscular immunizations

Mice were injected as previously described (46) with some modifications. 25 μg OVA (Grade V, Sigma) in 1 mg alum or alum alone was suspended in 100 μL 1X PBS. 50 μL per limb was injected in the right and left hind leg ventral muscles 14 and 21 days after *H. polygyrus bakeri* inoculation.

### Helminth infection

*Heligmosomoides polygyrus bakeri* (*H. polygyrus bakeri*) was propagated as previously described (47) and stored at 4°C until used. C57BL/6J mice were inoculated intragastrically with 200 third-stage larvae using a ball-tipped feeding needle. Adult worms in the intestinal contents were determined at sacrifice as previously described (47).

### OVA-TCR transgenic CD4^+^ T cell enrichment and adoptive transfer in helminth-infected and helminth-free mice

Spleens and MLN were harvested from OT-II (Thy1.1) mice and T lymphocytes were enriched using nylon wool fiber columns (Polysciences, Inc., Warrington, PA). CD4^+^ T cells were positively selected with CD4 (L3T4) magnetic microbeads (Miltenyi Biotec, Auburn, CA), pooled and suspended in PBS, and 4 to 6 x 10^6^ cells injected intravenously into C57BL/6 mice.

### Flow cytometric analysis

For the peripheral CD4^+^ to Foxp3^+^ Treg experiments, single-cell suspensions from MLN were prepared by passing tissue through a 70-μm cell strainer. For lamina propria (LP) cells, small intestinal segments were incubated in medium containing 3% FCS and 20 mM Hepes (HyClone) for 20 min at 37°C with continuous stirring. Tissue was then digested with 250 mg/ml liberase CI (Roche) and 500 mg/ml DNase I (Sigma-Aldrich), with continuous stirring at 37°C for 30 min. Digested tissue was forced through a Cellector tissue sieve, (Bellco Glass, Inc.) and strained through 70- and 40-μm cell strainers. To enrich for lymphocytes, the suspension was centrifuged at room temperature at 500 *g* for 20 min in 30% Percoll (GE Healthcare) in RPMI-1640. Cells were incubated with antibodies to Ly5.2 (clone 104), CD4 (clone RM4-5), CD25 (clone 7D4), CD103 (clone 2E7; all from eBioscience) and assessed for the expression of these markers in addition to eGFP by flow cytometry using an LSRII (BD Biosciences). Cells were also incubated with mAb against α_4_β_7_ (clone DATK32; BD Biosciences), CD44 (IM7; eBioscience), and 7-amino-actinomycin D (7-AAD; BD Biosciences) to detect dead cells. Cells were acquired with an LSR II flow cytometer (BD Biosciences) and flow cytometry data analyzed with FlowJo software (Tree Star, Ashland, OR).

For helminth-infection experiments, Thy1.1 FITC (clone OX-7), CD69 PE (clone H1.2F3), CD4 PerCP (clone RM4-5) and CD25 APC (clone PC61) and isotype controls were purchased from BD Biosciences. Non-specific binding was blocked with antibodies against CD16/CD32 (BD Biosciences, San Jose, CA). For intracellular cytokine staining MLN cells were stimulated as previously described (41) with some modifications. 2 x 10^6^ cells/ml were incubated for 24 h with 200 μg/ml ovalbumin protein (OVA, Grade V, Sigma, St.Louis, MO). Prior to being added to cultures, endotoxin levels in the OVA preparation were reduced to less than 0.7 EU/mg using a Detoxi-Gel endotoxin removal column (Pierce, Rockford, IL). During the final 4 h of culture, cells were pulsed with 12.5 ng/ml PMA (Sigma), 500 ng/ml ionomycin (Sigma), and 1 μg/ml GolgiPlug (BD Biosciences). Cells were harvested, surface stained and permeabilized with Cytofix/Cytoperm Buffer (BD Biosciences), washed with Perm/Wash Buffer (BD Biosciences) and stained with anti-IFN-γ APC (clone XMG1.2) and anti-IL-4 PE (clone 11B11) or anti-IL-10 APC (clone JES5-16E3), and anti-IL-13 PE (clone eBio13A, eBioscience, San Diego, CA). Cells were acquired using a FACScalibur (BD Biosciences) and data analyzed using FlowJo software (Tree Star, Ashland, OR).

### Measurement of serum and fecal antibody levels

Sera were collected weekly over the course of each experiment and feces were collected at sacrifice. Sera from individual mice were assayed for OVA-specific IgG1, IgG2b, IgG2c, and IgA by ELISA as previously described (27). For OVA-specific IgG1, IgG2b, and IgG2c, OD values were converted to ng/ml by comparison with a standard curve of anti-OVA Abs affinity purified from the serum of immunized C57BL/6 J mice using OVA conjugated to CNBr-activated Sepharose 4B (Amersham Biosciences, Uppsala, Sweden). To obtain ng/ml values of each anti-OVA Ab isotype, known amounts of purified mouse isotype control Abs from Southern Biotechnology Associates, Birmingham, AL (for IgG1 and IgG2b) or Bethyl Laboratories, Montgomery, TX (for IgG2c) were used. For OVA-specific IgA, OD values were converted to ng/ml of IgA by comparison with a purified IgA standard (BD Biosciences, San Jose, CA). Fecal extracts from individual mice were obtained as previously described (41) and OVA-specific IgA responses were determined by ELISA.

### Statistical analysis

Results are expressed as the mean ± standard error of the mean (SEM). One-way ANOVA followed by Tukey’s multiple comparisons test, unpaired t tests, or the Mann Whitney test were used to determine the significance of differences among helminth-free and helminth-infected vaccinated and unvaccinated groups of mice. Statistical differences were determined using GraphPad Prism (GraphPad Software, Inc., San Diego, CA). A *P* value of <0.05 was considered significant.

### Figure design

Figures were created using BioRender (https://biorender.com).

## RESULTS

### A live attenuated oral *Salmonella*-OVA vaccine disrupts antigen-specific regulatory networks to promote T-effector responses to OVA in the gut-associated lymphoid tissue

To determine the impact of oral vaccination with *Salmonella-*OVA on the development of OVA-specific regulatory T cells (Tregs) in the GALT, we introduced *Salmonella-*OVA into a congenic adoptive transfer model previously used to show that oral consumption of dietary OVA antigen drives conversion of OVA-specific T cells into Foxp3^+^ Tregs in the small intestinal lamina propria (LP) and gut-associated lymphoid tissue (GALT) (44). We adoptively transferred Ly5.2^+^ T cells from recombination-activating gene 1–deficient (RAG1 KO) OT-II transgenic (OT-II Tg) mice into Ly5.1^+^ RAG-1-replete B6.SJL recipients. Some recipients were then fed OVA antigen dissolved in drinking water (OVA-water) for five consecutive days. Others were orally vaccinated once with 10^12^ *Salmonella-*OVA or a sham live attenuated *Salmonella* vaccine strain that does not express OVA (*Salmonella*-BEM). Another subset received both OVA-water for five days and one dose of *Salmonella-*OVA (Fig. 1A). All CD4^+^ T cells in Ly5.2^+^ RAG-1 KO OT-II transgenic mice are specific for OVA and nearly all these CD4^+^ cells lack expression of the Treg transcription factor Foxp3 (Foxp3-expressing cells <0.05%) (44).We found that oral administration of OVA-water or one dose of *Salmonella-*OVA, but not the sham vaccine, increased the proportions of OVA-specific T cells in the GALT, particularly in the MLN (Fig. 1B and C). While OVA-specific T cells accumulated in the intestinal LP of mice fed OVA-water, whether or not they were vaccinated with *Salmonella-*OVA, oral vaccination with *Salmonella*-OVA induced a significantly lower frequency of OVA-specific T cells accumulating in the intestinal LP (Fig. 1D and E).

**Figure 1.**
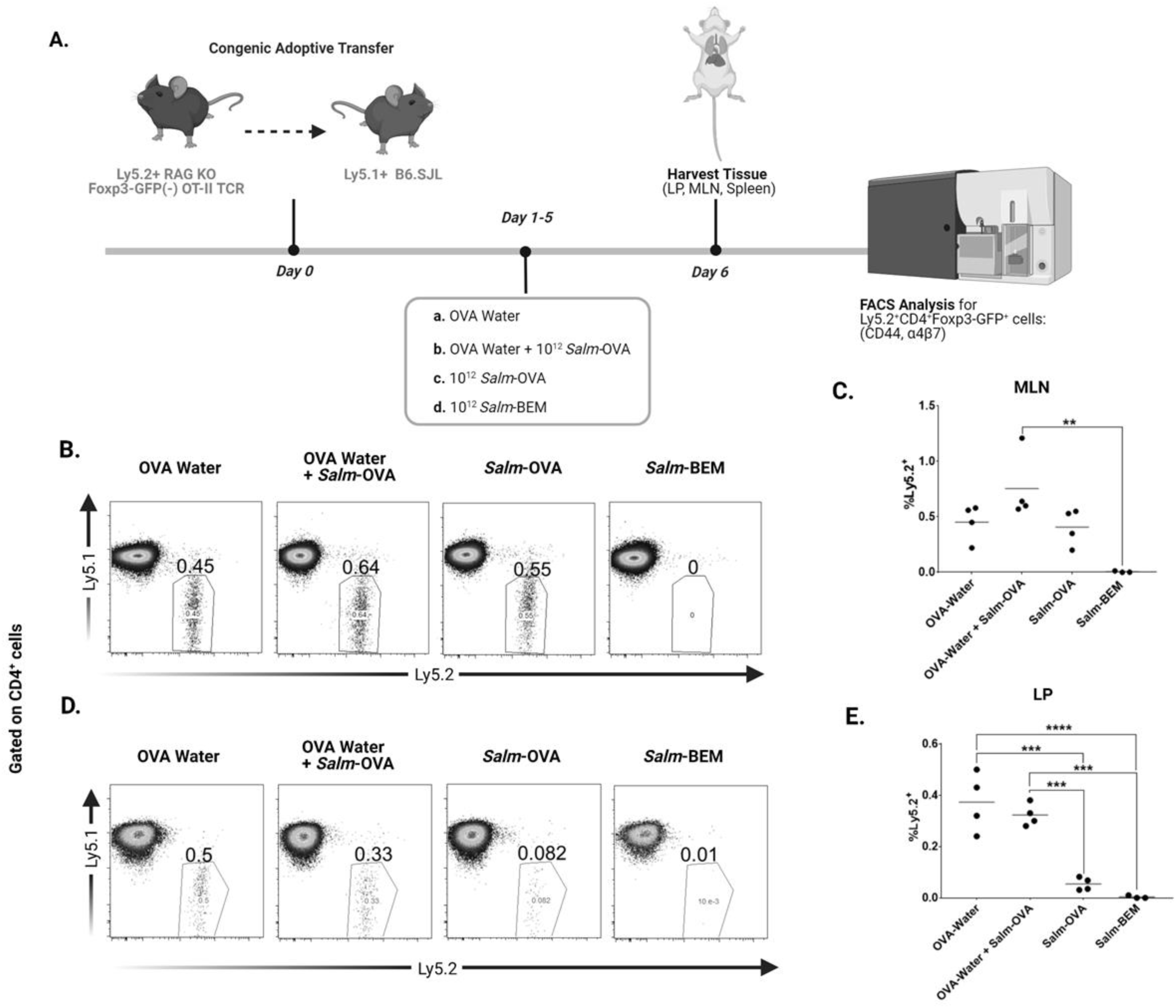
Feeding with soluble OVA antigen and/or oral vaccination with *Salmonella-*OVA induces accumulation of OVA-specific T cells in the GALT. (A) Experimental timeline. After gating on CD4^+^ T cells, transferred T cells in the MLN (B, C) or LP (D, E) of OVA antigen-fed mice or mice vaccinated with *Salmonella*-OVA (*Salm*-OVA) or the sham vaccine *Salmonella-* BEM (*Salm*-BEM) were identified by Ly5.2 expression. B and D are representative flow cytometry plots while C and E are summary graphs of the percentage of Ly5.2^+^ RAG1 KO OT-II T cells in MLN and LP respectively. Each dot in C and E represents a single mouse, with three to four mice per group. Experiment was repeated two times. Data shown are from one of two independent experiments. Statistical Analyses: One-way ANOVA followed by Tukey’s multiple comparisons test (\*\*\*\**P*<0.0001, \*\*\**P*<0.001, \*\**P*<0.01).

As expected, soluble OVA (OVA-water) induced the conversion of OVA-specific CD4^+^ T cells to Foxp3^+^ Tregs in MLN and intestinal LP ((44) and Fig. 2). However, oral vaccination with *Salmonella-*OVA impaired soluble OVA-driven conversion of OVA-specific CD4^+^ T cells to Foxp3^+^ Tregs in both the MLN and intestinal LP (Fig. 2). Moreover, oral vaccination with *Salmonella-*OVA increased the frequency of activated, OVA-specific, Foxp3^-^ effector T cells that expressed gut-homing surface molecules α4β7 and CD44 in both MLN (Fig. 3A-D) and LP (Fig 3E-H). *Salmonella-*OVA attenuated the increase in the frequency of α4β7- and CD44-expressing, OVA-specific, Foxp3^+^ Tregs in the MLN and LP normally induced by soluble OVA (Fig. 3). These data demonstrate that the GALT handles OVA expressed by a live attenuated oral *Salmonella* vaccine in a manner distinct from soluble OVA in drinking water and suggests that OVA acts as a vaccine antigen when introduced in the context of the *Salmonella-*OVA oral vaccine.

**Figure 2.**
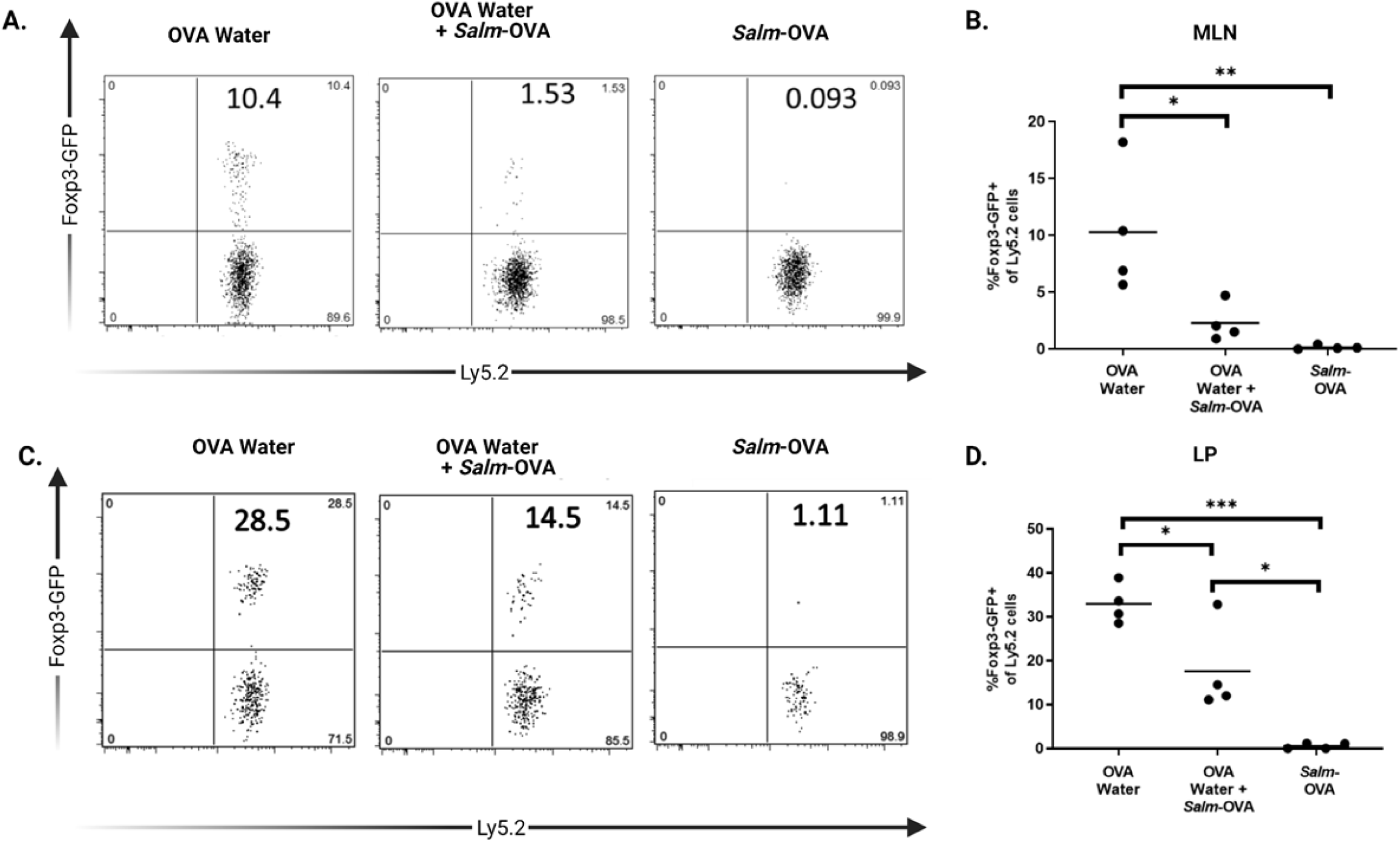
Oral vaccination with *Salmonella-*OVA disrupts oral soluble OVA-driven conversion of CD4^+^ T cells to Foxp3^+^ Tregs in the GALT. After gating on CD4^+^ T cells, transferred T cells in the MLN (A, B) or LP (C, D) of OVA antigen-fed or *Salmonella*-OVA (*Salm-*OVA) vaccinated mice were identified by Ly5.2 expression. Ly5.2^+^ cells were then assessed for intracellular Foxp3 expression. A and C are representative flow cytometry plots while B and D are summary graphs of the percentage of Foxp3^+^ among Ly5.2^+^ RAG1 KO OT-II T cells in MLN and LP respectively. Each dot in B and D represents a single mouse with four mice per group. Experiment was repeated two times. Data shown are from one of two independent experiments. Statistical Analyses: One-way ANOVA followed by Tukey’s multiple comparisons test (\*\*\**P*<0.001, \*\**P*<0.01, \**P*<0.05).

**Figure 3.**
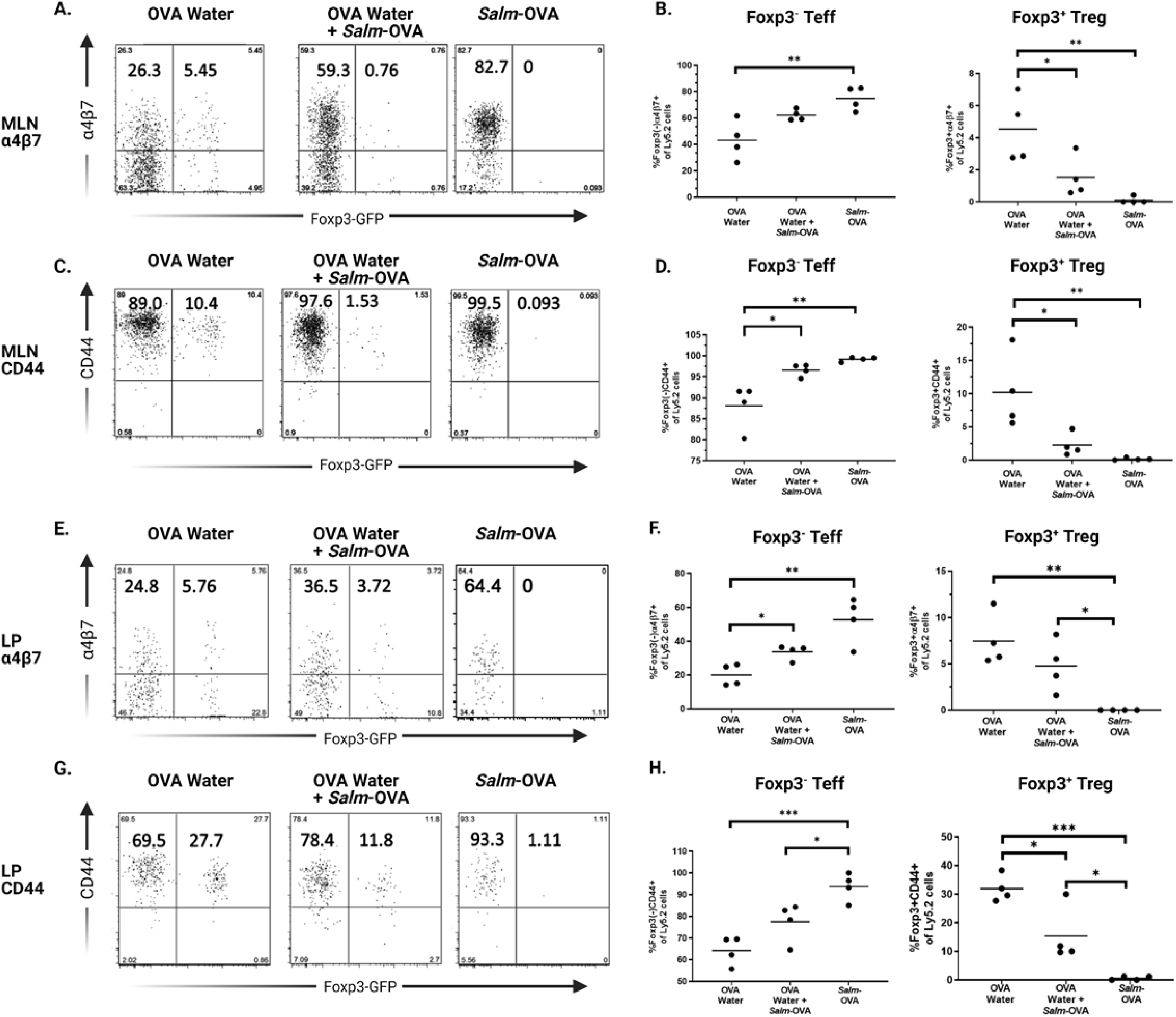
Oral vaccination with *Salmonella-*OVA increases the frequency of activated, OVA-specific Foxp3^-^ T effectors and decreases the frequency of OVA-specific Foxp3^+^ Tregs expressing gut homing molecules in the MLN and lamina propria. After gating on CD4^+^ T cells, transferred T cells in the MLN of OVA antigen-fed or *Salmonella*-OVA vaccinated mice were identified by Ly5.2 expression. Ly5.2^+^ cells in the MLN (A-D) and LP (E-H) were then assessed for intracellular Foxp3 expression and cell surface expression of α4β7 (A, B, E, F) or CD44 (C, D, G, H). A, C, E, and G are representative flow cytometry plots while B, D, F and H are summary graphs. Each dot represents a single mouse with four mice per group. Experiment was repeated two times. Data shown are from one of two independent experiments. Statistical Analyses: one-way ANOVA followed by Tukey’s multiple comparisons test (\*\*\**P*<0.001, \*\**P*<0.01, \**P*<0.05).

### Preexisting intestinal helminth infection reduces Th1-skewed OVA-specific antibody responses to oral vaccination with *Salmonella-*OVA

Chronic helminth infection of at least two weeks duration results in significant impairment of host immune responses to Th1/IFN-γ inducing malaria infection (26), IL-12 /IFN-γ dependent trinitrobenzenesulfonic acid (TNBS)-induced colitis (48), and parenteral vaccination against yellow fever virus YFV-17D (22). To determine whether chronic intestinal helminth infection could alter antibody responses to oral immunization, C57BL/6 mice were infected or not with the natural murine gastrointestinal helminth, *H. polygyrus bakeri* 14 days prior to oral vaccination with *Salmonella-*OVA or the sham vaccine *Salmonella*-BEM (Fig. 4A). We examined OVA-specific antibody responses to *Salmonella-*OVA or sham vaccine in sera and fecal extracts of helminth-infected and helminth-free mice (Fig. 4B-F). As expected, mice vaccinated with *Salmonella-*BEM did not make OVA-specific antibody responses ((41) and Fig. 4). Helminth-free mice vaccinated with *Salmonella-*OVA made a highly Th1-skewed OVA-specific serum IgG2c response, 10 to 15-fold greater than the OVA-specific serum IgG2b and IgG1 responses, respectively (Fig. 4B-D and (41)). OVA-specific IgG2b and IgG2c levels were significantly lower in vaccinated helminth-infected mice compared to vaccinated helminth-free mice (Fig. 4C, D). Helminth infection delayed but did not eliminate the OVA-specific serum IgA response to *Salmonella-*OVA (Fig. 4E). OVA-specific fecal IgA responses were reduced two-fold in helminth-infected vaccinated mice compared to helminth-free mice although this did not reach statistical significance (Fig. 4F).

**Figure 4.**
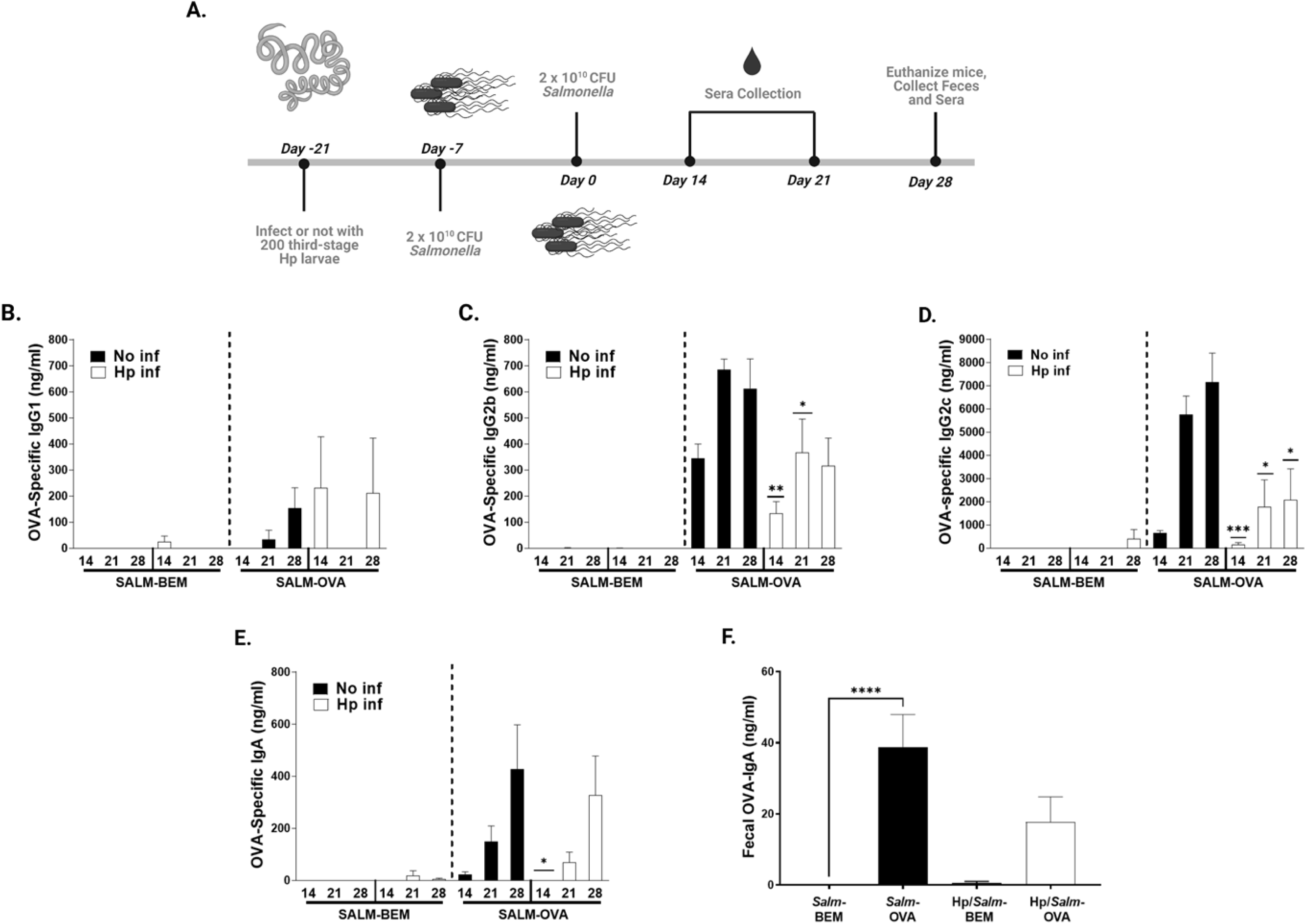
Th1-skewed antibody responses to oral vaccination with *Salmonella-*OVA are reduced in mice with preexisting helminth infection. (A) Experimental timeline; 14 days after intragastric inoculation with 200 third-stage *H. polygyrus bakeri* larvae, helminth-free (B-E; No Inf, black bars) and helminth-infected (B-E; Hp inf, white bars) C57BL/6 mice were given two intragastric doses of 2 x 10^10^ *Salmonella*-BEM (SALM-BEM) or *Salmonella-*OVA (SALM-OVA) and OVA-specific serum (B) IgG1, (C) IgG2b, (D) IgG2c, and (E) IgA were measured 14, 21, and 28 days post vaccination by ELISA. OVA-specific IgA in fecal extracts was measured 28 days post vaccination by ELISA (F). Pooled data from two independent experiments (n=11-12 mice per group). Statistical Analyses: Unpaired t-test in B-E comparing helminth-infected to helminth-free at same time point; One-way ANOVA in F followed by Tukey’s multiple comparisons test. \*\*\**P*<0.001, \*\**P*<0.01, \**P*<0.05.

Preexisting helminth infection did not enhance OVA-specific IgG1 responses in orally vaccinated mice (Fig. 4B) despite the Th2 polarized helminth-induced polyclonal serum and fecal IgG1 and serum IgE responses in helminth-infected mice (Supplementary Fig. 1B-D). We found significantly elevated levels of total serum IgE in helminth-infected mice vaccinated with *Salmonella* compared to their unvaccinated, helminth-infected counterparts, perhaps due to enhanced polyclonal B cell activation in the presence of *Salmonella* LPS, as reported in *in vitro* studies by others (49). The mean number of parasites recovered at sacrifice from the intestinal contents of mice given the sham vaccine *Salmonella*-BEM was lower than that recovered from unvaccinated helminth-infected mice, although this did not reach statistical significance (Supplementary Fig. 1E). There was no difference in mean number of parasites recovered from mice vaccinated with *Salmonella-*OVA compared to unvaccinated helminth-infected mice (Supplementary Fig. 1E).

### Intestinal helminth infection reduces OVA-specific IgG responses to intramuscular vaccination with OVA and the non-microbial adjuvant alum

To determine whether intestinal helminth infection could suppress vaccine antigen-specific antibody responses to a parenterally-administered model OVA protein subunit vaccine, we inoculated C57BL/6 mice with helminth 14 days prior to intramuscular (i.m.) vaccination with OVA adsorbed to the vaccine adjuvant alum (OVA-alum, Fig. 5A). We observed that helminth-free mice vaccinated i.m. with OVA-alum made a highly Th2-skewed OVA-specific serum IgG1 response (Fig. 5B). Th1-dependent OVA-specific serum IgG2c was not detected and OVA-specific IgG2b levels were 150-fold lower than the OVA-IgG1 levels (data not shown). Notably, helminth-infected mice vaccinated i.m. with OVA-alum made significantly lower OVA-specific IgG1 (Fig. 5B) and IgG2b (data not shown) responses when compared to helminth-free vaccinated mice despite elevated, Th2-skewed, polyclonal serum IgG1 and IgE levels (Fig. 5C, D). Comparable numbers of adult worms could be recovered from the intestinal contents of both vaccinated and unvaccinated helminth-infected mice (Fig. 5E). Th2-skewed antibody responses to i.m. OVA-alum vaccination remained significantly reduced in helminth-infected mice, despite robust polyclonal IgG1 and IgE responses associated with helminth infection, even when mice were housed in a specific pathogen free facility in a completely different institution than in Fig. 5 (see Supplementary Fig. 2). Thus, chronic intestinal helminth infection impaired immune responses to vaccines delivered via either mucosal or parenteral routes.

**Figure 5.**
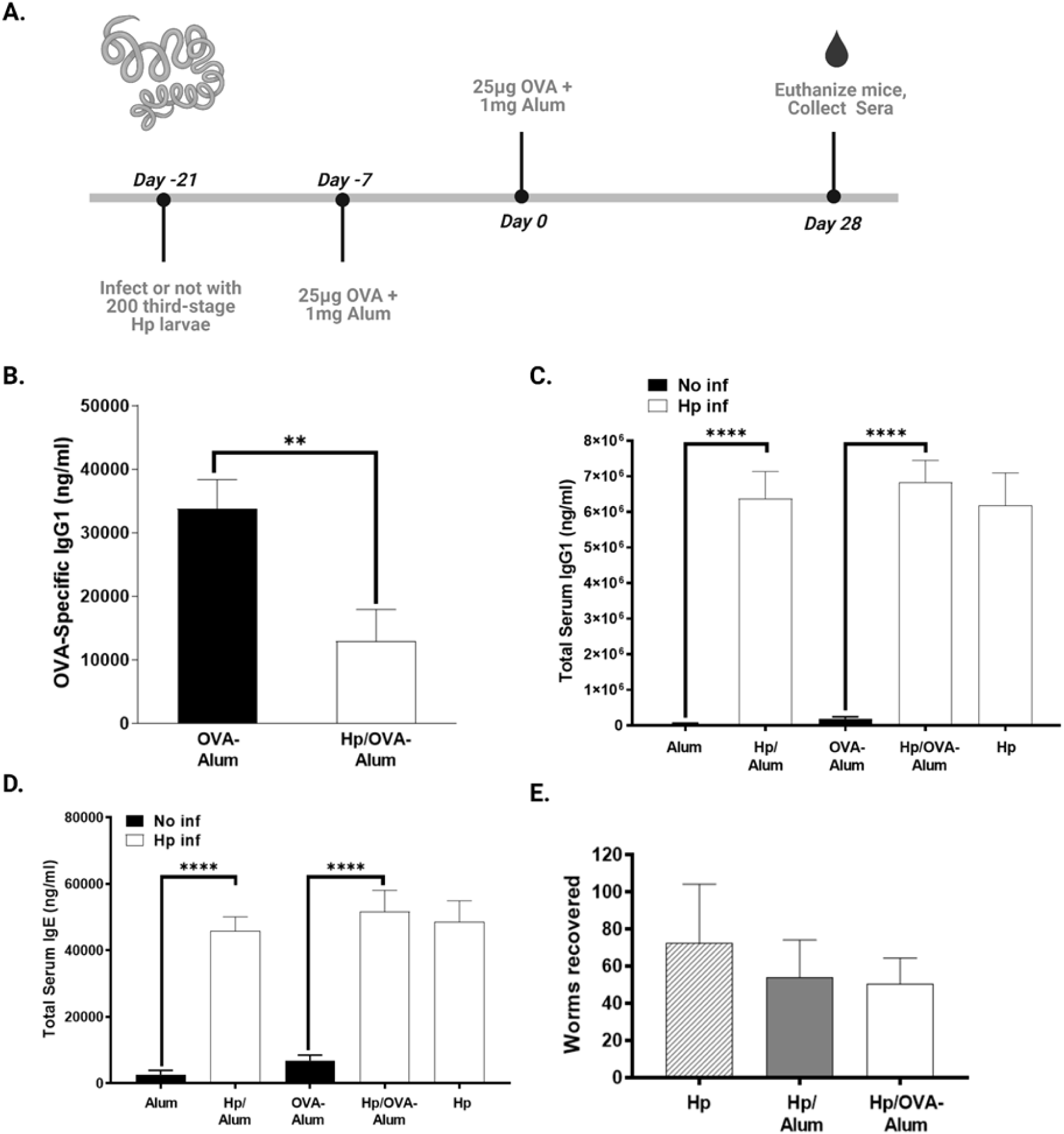
Th2-skewed antibody responses to intramuscular vaccination with OVA-alum are significantly reduced while polyclonal IgG1 and IgE responses are elevated in vaccinated, helminth-infected mice. (A) Experimental timeline; 14 days after intragastric inoculation with 200 third-stage *H. polygyrus bakeri* larvae, helminth-free C57BL/6 mice (black bars) and helminth-infected C57BL/6 mice (white bars), were given two intramuscular doses of 25 mg OVA adsorbed to 1 mg alum or 1mg alum alone spaced one week apart. (B) OVA-specific IgG1. (C) Total Serum IgG1. (D) Total serum IgE. (E) Worms recovered. Data in B-E are pooled from two independent experiments; n=8-10 mice per group. Statistical Analyses: unpaired t-test in B; One-way ANOVA followed by Tukey’s multiple comparisons test in C-E (\*\*\*\**P*<0.0001, \*\**P*<0.01).

### Helminth infection does not reduce splenic bacterial titers and oral *Salmonella* does not alter helminth-induced organomegaly

Since SL3261, the parent strain of *Salmonella-*BEM and *Salmonella-*OVA, is highly attenuated, its ability to replicate *in vivo* is limited; however, after intragastric administration, bacteria disseminate systemically and are recoverable from the spleens three days after vaccination and up until four weeks later (data not shown). To determine whether helminth-mediated suppression of antibody responses to oral *Salmonella-*OVA was due to alterations in the systemic dissemination of the vaccine, we examined bacterial titers in the spleens of helminth-infected and uninfected mice three days after oral vaccination (Supplementary Fig. 3A). We found no difference in CFU per gram tissue recovered from the spleens of helminth-infected and helminth-free mice vaccinated with *Salmonella-*OVA or *Salmonella-*BEM (Supplementary Fig. 3B), suggesting that intestinal helminth infection did not alter systemic trafficking of the live attenuated vaccines. Conversely, ten days after *H. polygyrus bakeri* infection (three days after oral vaccination), comparable numbers of adult worms could be recovered from the intestinal contents of both vaccinated and unvaccinated mice (Supplementary Fig. 3C). Both the draining MLN and spleens of helminth-infected mice were enlarged compared to helminth-free mice and significantly greater in mass, regardless of whether the mice were vaccinated with *Salmonella-*BEM or *Salmonella-*OVA (Supplementary Fig. 3D, E). Taken together, these data suggest that helminth infection does not impair systemic spread of the *Salmonella* vaccines.

### Activated vaccine antigen-specific CD4^+^ T cells accumulate in the draining MLN of both helminth-free and helminth-infected mice vaccinated with *Salmonella-*OVA

Cytokines produced by antigen-activated CD4^+^ helper T cells typically drive antibody class switching and stimulate B cells to produce antibodies against T-cell dependent protein antigens (50). To determine if impaired antigen-specific humoral responses in helminth-infected mice were due to a defect in the response to vaccine antigen by antigen-specific CD4^+^ T helper cells, we adoptively transferred C57BL/6 mice (whose T cells express the surface marker Thy1.2) with CD4^+^Thy1.1^+^ OVA-specific T cell receptor transgenic OT-II cells. Two days later, a subset of mice were infected with helminth larvae. Following helminth infection, mice were orally vaccinated with either *Salmonella-*OVA or the sham vaccine *Salmonella-*BEM (Fig. 6A). Three days after vaccination, both the frequency and total number of OVA-specific CD4^+^Thy1.1^+^ OT-II cells in the MLN were higher in helminth-free and helminth-infected mice vaccinated with *Salmonella-*OVA compared to *Salmonella-*BEM vaccinated mice (Fig. 6B-D). The mean frequency, but not mean total number, of MLN OT-II cells was significantly lower in helminth-infected, *Salmonella-*OVA vaccinated mice than in their uninfected, *Salmonella-*OVA vaccinated counterparts (Fig. 6C, D). This was likely due to a helminth-induced influx of helminth-specific effector cells into the MLN, reflected in the larger organ mass in helminth-infected vaccinated mice (Supplementary Fig. 3D) and increased total cell numbers in the draining MLN of helminth-infected mice (data not shown). However, the proportion and total number of OT-II cells that expressed CD69, a marker of early lymphocyte activation, were higher in both helminth-free and helminth-infected mice vaccinated with *Salmonella-*OVA compared to mice vaccinated with *Salmonella-*BEM (Fig. 6E, F).

**Figure 6.**
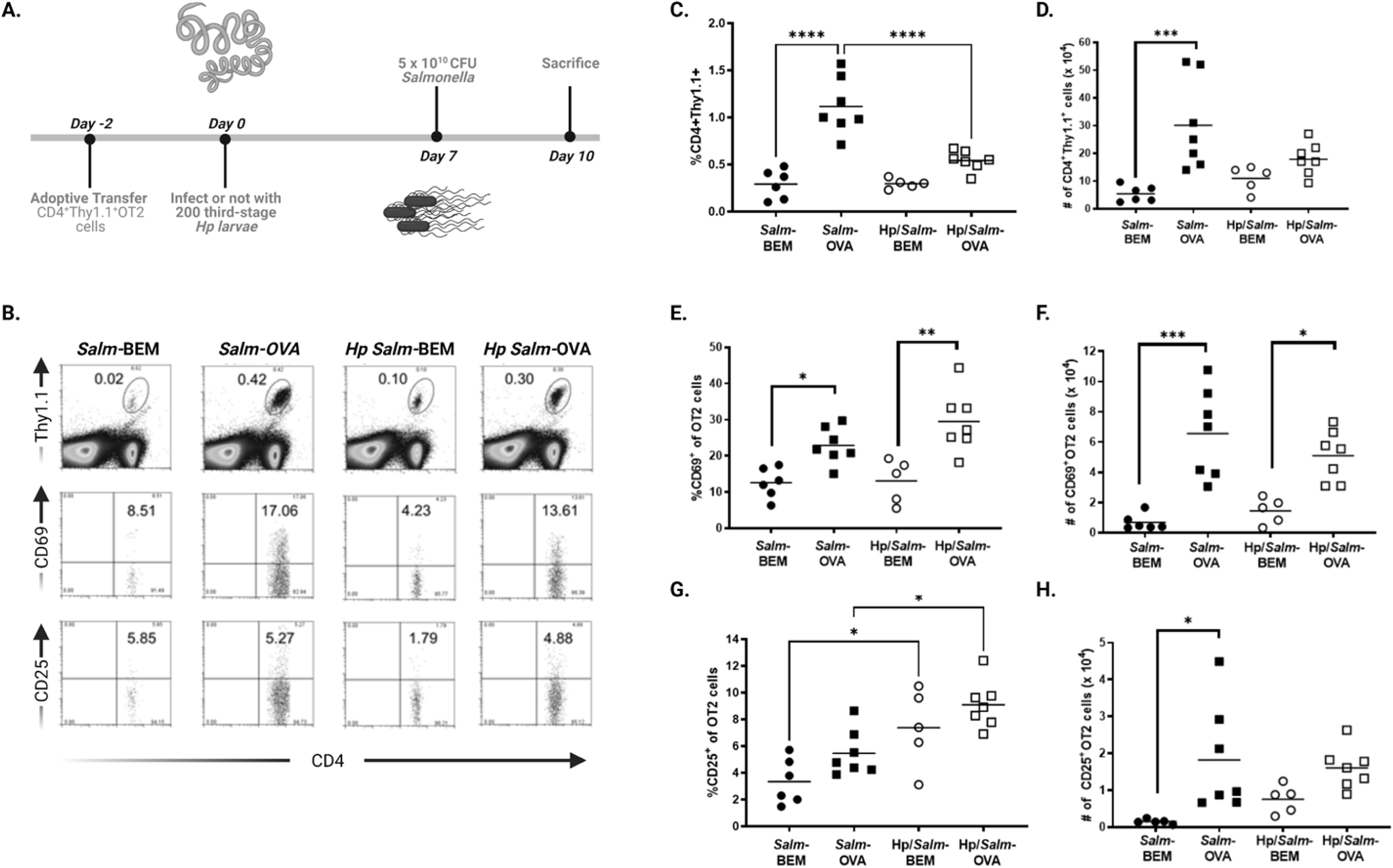
Activated vaccine antigen-specific CD4^+^ T cells accumulate in the draining MLN of both helminth-free and helminth-infected mice vaccinated with *Salmonella-*OVA. (A) Experimental timeline. (B) Representative flow cytometry plots showing adoptively transferred CD4^+^Thy1.1^+^ OVA-TCR transgenic OT-II cells, percent CD69^+^ and percent CD25^+^ among OT-II cells in MLNs of helminth-free and helminth-infected mice. (C) Proportion of adoptively transferred CD4^+^Thy1.1^+^ OVA-TCR transgenic OT-II cells in MLNs of helminth-free (black symbols) and helminth-infected (white symbols) mice. (D) Total number OT-II cells recovered from MLN. (E) Percent CD69^+^ among OT-II cells. (F) Total number CD69^+^OT-II cells (G) Percent CD25^+^ among OT-II cells (H) Total number CD25^+^OT-II cells. Salm-BEM (circles) = *Salmonella-*BEM. Salm-OVA (squares) = *Salmonella-*OVA. Hp = *H. polygyrus bakeri*. Symbols represent individual mice; lines represent mean percentages. Pooled data from three independent experiments; n=5 to 7 mice per group. Statistical Analyses: one-way ANOVA followed by Tukey’s multiple comparisons test (\*\*\*\**P*<0.0001, \*\*\**P*<0.001, \*\**P*<0.01, \**P*<0.05).

Both activated T-effector and Treg populations can express IL-2Rα chain, CD25 (51). We found a modest increase in the percentage, and a significant increase in the number, of CD25^+^ OT-II cells found in MLNs from helminth-free mice vaccinated with *Salmonella-*OVA compared to helminth-free mice that received the sham vaccine (Fig. 6G, H). In addition, the percentage of CD25^+^ OT-II cells in helminth-infected, vaccinated mice was nearly 2-fold greater than in helminth-free mice (Fig. 6G). However, there was no significant difference in total number of CD25^+^ OT-II cells recovered from the MLNs of helminth-infected mice vaccinated with *Salmonella*-OVA compared to helminth-free mice (Fig. 6H). By contrast, total numbers of MLN, non-TCR transgenic, CD4^+^Thy1.1^-^CD25^+^ cells were 2-fold greater in helminth-infected, vaccinated mice than in helminth-free mice (Supplementary Fig. 4), suggesting that helminth infection increased total numbers of activated effector and regulatory T cells in orally vaccinated mice.

### Helminth-induced Th2-polarized cytokine responses are intact in orally vaccinated mice

Intestinal helminth infection promotes the production of Th2 effector cytokines, including IL-4 and IL-13 (52), and regulatory cytokines like IL-10 and TGF-β (53) by CD4^+^ T cells. We examined cytokine responses in polyclonal and OVA-specific CD4^+^ T cell populations following oral vaccination in helminth-free and helminth-infected mice using intracellular staining and flow cytometry (Fig. 7). Ten days after helminth infection and three days after oral vaccination, we found comparable frequencies of IFN-γ^+^CD4^+^Thy1.1^-^ non-TCR transgenic Th1 cells in the draining MLN among helminth-infected and helminth-free mice (Fig. 7B and C). However, a significantly greater percentage of CD4^+^Thy1.1^-^ cells were IL-4^+^ (Fig. 7D), IL-10^+^ (Fig. 7E), and IL-13^+^ (Fig. 7F) in both vaccinated and unvaccinated, helminth-infected mice compared to uninfected mice. Thus, oral vaccination with the Th1-polarizing attenuated *Salmonella* vaccine did not prevent the generation of a robust cell-mediated Th2 and Treg cytokine response to helminth infection.

**Figure 7.**
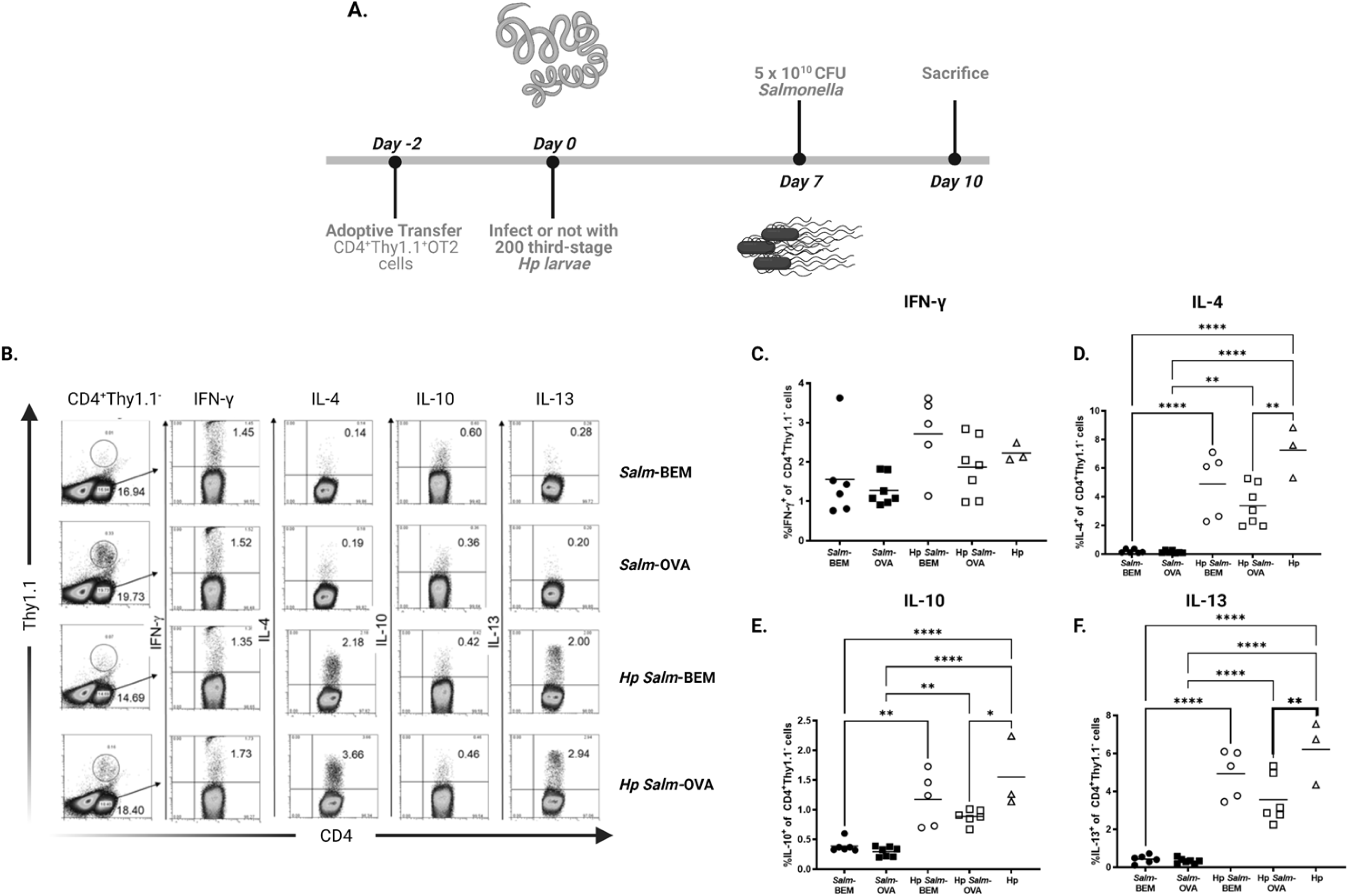
Intestinal helminth infection induces Th2-polarized cytokine responses in CD4^+^ T cell populations in both vaccinated and unvaccinated mice. (A) Experimental timeline; two days after adoptive transfer of 4 to 6 x 10^6^ CD4^+^Thy1.1^+^ OVA-TCR transgenic OT-II cells, mice were infected (white symbols) or not (black symbols) with 200 third-stage *H. polygyrus bakeri* (Hp) larvae. Seven days after helminth infection, mice received one intragastric dose of ∼5 x 10^10^ *Salmonella-*BEM or *Salmonella-*OVA. 3 days later, MLN cells were harvested, cultured overnight with OVA, pulsed for 4 h with PMA, ionomycin, and Golgiplug and surface labeled with mAbs to CD4 and Thy1.1, fixed, permeabilized and intracellularly stained with Abs against IFN-γ, IL-4, IL-10, and IL-13. (B) Representative flow cytometry plots and summary graphs showing (C) percent IFN-γ^+^ (D) percent IL-4^+^ (E) percent IL-10^+^ (F) percent IL-13^+^ among non-TCR transgenic CD4^+^Thy1.1^-^ MLN cells. Salm-BEM (circles) = *Salmonella-*BEM. Salm-OVA (squares) = *Salmonella-*OVA. Hp (triangles) = *H. polygyrus bakeri* only. Symbols represent individual mice; lines represent mean percentages. Pooled data from 3 independent experiments; n=3 to 7 mice per group. Statistical Analyses: One-way ANOVA followed by Tukey’s multiple comparisons test (\*\*\*\**P*<0.0001, \*\*\**P*<0.001, \*\**P*<0.01, \**P*<0.05).

### Vaccine antigen-specific CD4^+^ T cells from the draining MLN of helminth-infected, orally vaccinated mice produce Th2-type effector and regulatory cytokines

We next examined Th1 (IFN-γ), Th2 (IL-4, IL-13), and Treg (IL-10) cytokines in OVA-specific CD4^+^Thy1.1^+^ OT-II MLN cells re-stimulated with OVA *in vitro* (Fig. 8). The frequencies and total numbers of OT-II cells recovered from cultured MLN cells of helminth-free and helminth-infected *Salmonella-*OVA vaccinated mice were significantly greater than in helminth-free, *Salmonella-*BEM vaccinated mice (Fig. 8C, D). Although the percentage and total numbers of IL-13^+^ and IFN-γ^+^OT-II cells were not statistically significantly different between helminth-free and helminth-infected vaccinated mice, the percentage and total numbers of OT-II cells producing, IL-4, and IL-10 was significantly increased in helminth-infected, vaccinated mice compared to uninfected, vaccinated mice (Fig. 8E-L). The helminth-modified Th2 response is characterized by elevated antigen-specific Th2-type cytokine production in conjunction with elevated antigen-specific IL-10 production to heterologous antigens administered to helminth-infected mice (54). The increased frequency and total number of Th2-type IL-4^+^ OT-II cells coupled with enhanced percentages and total numbers of OVA-specific cells producing IL-10 was consistent with the development of a helminth-modified Th2 and Treg response to oral vaccination with *Salmonella-*OVA. The increased frequency and total number of OVA-specific CD4^+^ IL-10-producing T cells and the drop in serum antibody responses to OVA in helminth-infected, vaccinated mice suggests a role for IL-10-secreting CD4^+^ T regulatory cells in reducing vaccine-induced humoral responses in helminth-infected mice.

**Figure 8.**
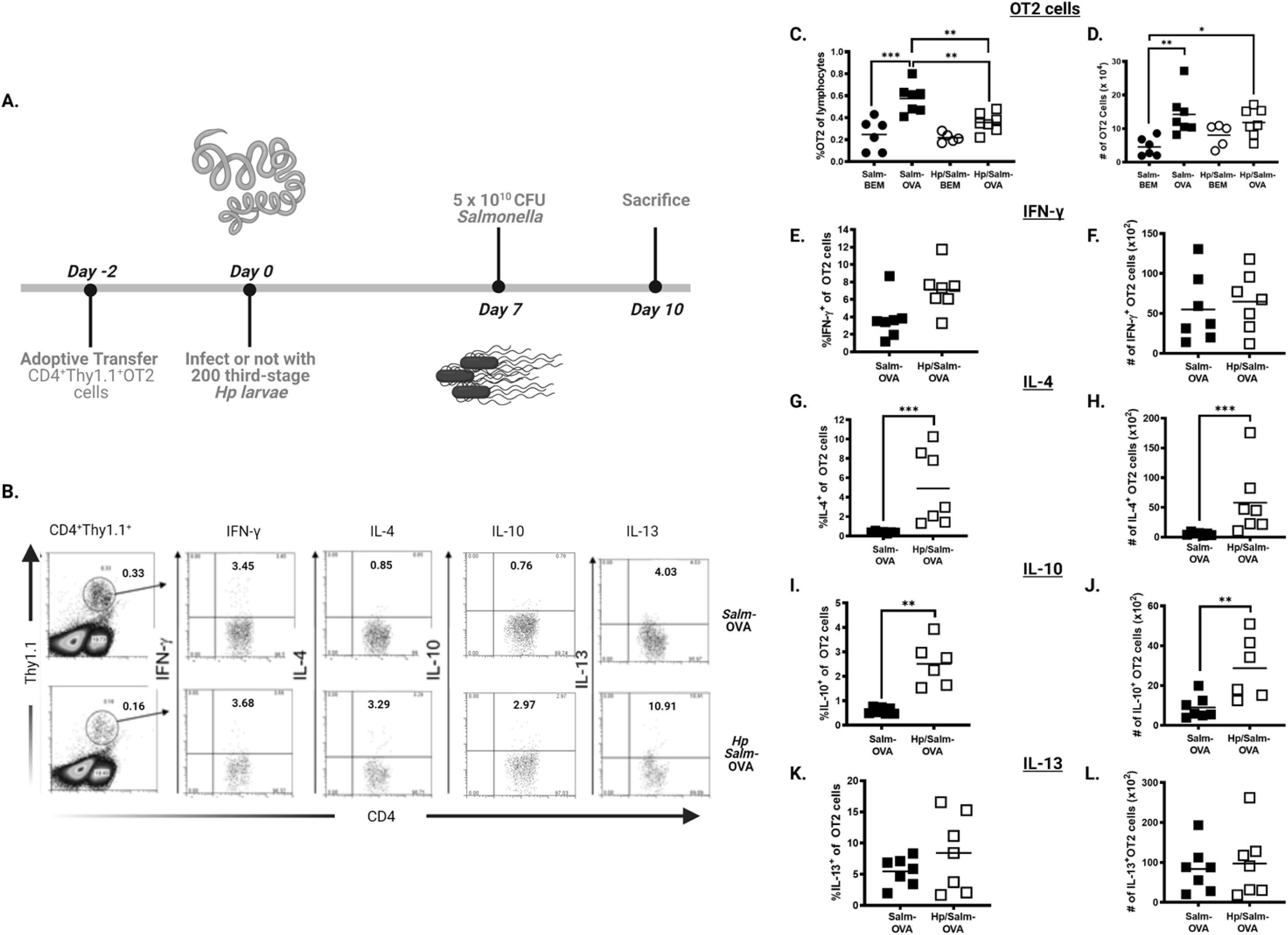
OVA-specific CD4^+^ T cells from the draining MLN of helminth-infected mice vaccinated with *Salmonella-*OVA produce Th2 effector and regulatory cytokines. (A) Experimental timeline. (B) Representative flow cytometry plot and (C-L) summary graphs showing (C) percent CD4^+^Thy1.1^+^ and (D) total number of CD4^+^Thy1.1^+^ among re-stimulated MLN cells (E) percent IFN-γ^+^ and (F) total number of IFN-γ^+^ (G) Percent IL-4^+^ (H) total number IL-4^+^ (I) percent IL-10^+^ and (J) total number IL-10^+^ (K) percent IL-13^+^ and (L) total number IL-13^+^ of adoptively transferred CD4^+^Thy1.1^+^ OT-II MLN cells. MLN cells were harvested, cultured overnight with OVA, pulsed for 4 h with PMA, ionomycin, and Golgiplug and surface labeled with mAbs to CD4 and Thy1.1, fixed, permeabilized and intracellularly stained with Abs against IFN-γ, IL-4, IL-10, and IL-13. Helminth-free (black symbols); helminth-infected (white symbols). Salm-BEM (circles) = *Salmonella-*BEM. Salm-OVA (squares) = *Salmonella-*OVA. Symbols represent individual mice; lines represent mean percentages. Pooled data from three independent experiments; n=5 to 7 mice per group. Statistical Analyses: one-way ANOVA followed by Tukey’s multiple comparisons test (C, D) and Mann Whitney test (E-L. \*\*\**P*<0.001, \*\**P*<0.01, \**P*<0.05).

## DISCUSSION

In humans and mice, protective immune responses against oral *Salmonella* vaccines involve both T and B cell responses, including robust expansion of CD4^+^ Th1 cells, IFN-γ production (17, 18), and *Salmonella*-specific IgG and IgA responses that facilitate clearance of the organism (18, 19). We and others have shown that the host immune response to heterologous vaccine antigen produced within the recombinant attenuated oral *Salmonella* vaccine (RASV) system is a CD4^+^ Th1-biased immune response (41, 55) that depends on intact signaling via MyD88 (41). The experiments presented here expand our current understanding of how the RASV system induces immunity to heterologous vaccine antigens. We show that the RASV system disrupts vaccine antigen-specific regulatory T cell networks in the gut-associated lymphoid tissue (GALT). It reduces the frequency of activated, vaccine-antigen specific, Foxp3^+^ regulatory T cells in the GALT that express gut-homing markers. The RASV system concurrently promotes the accumulation of activated, vaccine antigen-specific, Foxp3^-^ effector T cells expressing gut-homing surface molecules in the GALT (Figs. 2 and 3).

Because they are versatile and reliably induce Th1-biased immune responses, live attenuated oral *Salmonella* vaccine strains are widely used in agriculture, veterinary medicine, and preventative care of humans to protect against salmonellosis (11, 12), typhoid (13, 14) and paratyphoid fever (15). The recombinant attenuated oral *Salmonella* vaccine (RASV) system has also been used as an experimental vaccine platform to develop oral vaccine candidates for protection against food borne parasites (56, 57), human papilloma virus (58), streptococcal pneumonia (55), and shigellosis (59), among many other pathogens. Yet, disparities in the immunogenicity of oral *Salmonella* vaccines (60, 61) and other oral and parenteral vaccines in low- and middle-income countries compared to high-income countries have been repeatedly described (62–65).

One compelling hypothesis for this disparity is that endemic helminth infection alters immune responses to vaccination, and indeed, in human population studies, multiple reports describe decreased vaccine efficacy in people with chronic helminth infections (60, 66–69). In this study, we used a murine model to examine the impact of intestinal helminth infection on the response to vaccination. We demonstrated that chronic infection with the intestinal helminth *Heligmosomoides polygyrus bakeri* significantly suppressed Th1-skewed OVA-specific antibody responses to our live attenuated oral *Salmonella*-OVA vaccine (Fig. 4). Strikingly, despite robust helminth-induced Th2-biased total IgG1 and IgE responses in helminth-infected, vaccinated mice (Supplementary Fig. 1), *H. polygyrus bakeri* infection failed to enhance the development of a Th2-dependent OVA-specific IgG1 response to *Salmonella-*OVA (Fig. 4).

The reduced antibody responses to oral vaccination in helminth*-*infected mice were not due to an impaired ability of the live attenuated *Salmonella* to traffic systemically and reach immune organs like the spleen. By day 3 after oral vaccination, comparable CFU *Salmonella* per gram tissue were recoverable from the spleens of helminth-free and helminth-infected mice (Supplementary Fig. 3). The reduced antigen-specific humoral responses were also not due to impaired ability of adaptive immune cells in helminth-infected mice to recognize and respond to vaccine antigens. OVA-specific CD4^+^ T cells expressing the activation marker CD69 accumulated in the draining MLN of both helminth-free and helminth-infected mice vaccinated with the OVA-expressing *Salmonella* (Fig. 6). Moreover, similar numbers of OVA-specific CD4^+^ T cells in helminth-infected and helminth-free mice vaccinated with *Salmonella-*OVA produced the Th1 effector cytokine IFN-γ when re-stimulated *in vitro* with OVA (Fig. 8).

Vaccination with *Salmonella*-OVA did not alter helminth-induced organomegaly (Supplementary Fig. 3) nor did it hinder the development of Th2-polarized cytokine responses in CD4^+^ T cells from vaccinated mice (Fig. 7). Notably, helminth infection primed for a Th2-biased and Treg-biased cytokine response to an ordinarily Th1-biasing vaccine, inducing greater frequencies of IL-4 and IL-10-producing vaccine antigen-specific CD4^+^ T cells (Fig. 8). The elevation in Th2 and Treg cytokines that we observed, in both polyclonal and vaccine antigen-specific T cell populations, mirrors the helminth-modified Th2 response to heterologous antigens previously reported by Mangan and colleagues in a mouse model of allergen-induced airway disease with concomitant helminth infection (54). This signature cytokine pattern has been observed in a variety of allergic and inflammatory disease models in our lab and others (29, 48). Helminth-induced IL-10 production in particular has been implicated in protecting against both chemically-induced, colonic inflammation (48, 70) and allergic inflammation (29, 71). While the helminth-modified, Th2 cytokine response is beneficial and protective in these inflammatory disease models, our data suggest that it is detrimental for generating robust immune responses in our helminth infection/vaccine model. IL-10-secreting, CD4^+^ T cell populations are associated with helminth-mediated immune suppression, as are CD4^+^CD25^+^ T cells, even in the absence of IL-10 secretion (72). Accordingly, we found an increased frequency of polyclonal (Fig. 7) and vaccine-antigen specific (Fig. 8) CD4^+^ IL-10-producing T cells and higher numbers of polyclonal CD4^+^CD25^+^ T cells (Supplementary Fig. 4) in helminth-infected compared to helminth-free mice.

Helminth-induced alterations in vaccine antigen-induced cytokine production have been previously described in both human studies and mouse models (31, 67, 69, 73). Elias et al. observed reduced purified protein derivative (PPD)-specific IFN-γ secretion by peripheral blood mononuclear cells isolated from bacilli Calmette-Guerin (BCG)-vaccinated Ethiopian subjects with concomitant intestinal helminth infection when compared to anthelmintic-treated controls (69). Su et al. and Nookala et al. both reported decreased vaccine antigen-induced IFN-γ in their models, with enhanced production of *P. chabaudi* antigen-specific IL-4, IL-13, and IL-10 in an intestinal helminth infection/malaria vaccination model (31) and enhanced tetanus toxoid-specific IL-10 in the human lymphatic filariasis/tetanus vaccination study (73). Surprisingly, we found no difference in the frequencies of polyclonal IFN-γ^+^CD4^+^ T cells between helminth-free and helminth-infected mice (Fig. 7). There were also comparable frequencies and total numbers of vaccine antigen-specific IFN-γ^+^ CD4^+^ T cells in helminth-infected and helminth-free vaccinated mice (Fig. 8). This may reflect the potent Th1-inducing properties of our live attenuated *Salmonella* vaccine compared to the protein subunit plus adjuvant vaccines employed in the other studies.

Intestinal helminth infection induced OVA-specific, Th2 cytokine-producing CD4^+^ T cells after oral *Salmonella-*OVA vaccination, but this did not translate into enhanced vaccine antigen-specific, Th2-dependent IgG1 antibody production in helminth-infected mice. Even in the context of intramuscular vaccination with OVA adsorbed to the adjuvant alum, which promotes Th2-skewed antibody responses to co-administered protein antigens (74, 75), intestinal helminth infection suppressed Th2-dependent, antigen-specific IgG1 production (Fig. 5 and Supplementary Fig. 2). Intestinal helminth infection has been shown to modulate cellular immune responses to i.m. and intravenous vaccination in mice (31, 32). Chronic co-infection with multiple viral pathogens in conjunction with the intestinal helminth *H. polygyrus bakeri* can reduce serum antibody responses to subcutaneous injection of the yellow fever vaccine (22). Our study highlights the suppressive effect of intestinal helminth infection on vaccine antigen-specific antibody responses to i.m. vaccination, even in the absence of any other infection.

The discordance of the impact of helminth-infection on vaccine-induced, T-effector cell responses compared to humoral immune responses (76) may depend on worm burden and chronicity of helminth infection, as has been shown in epidemiologic studies examining the effects of helminth infection on allergic disease (33). Individuals with chronic helminth infection and heavy worm burdens in a Venezuelan study were protected from atopic skin reactivity against house dust mite antigen, whereas those with sporadic infection and light worm burdens had elevated allergen-specific IgE responses and high skin reactivity (77). Su et al. have reported a similar phenomenon in their malaria/intestinal helminth coinfection model; while *H. polygyrus bakeri* infection of one week duration could suppress antimalarial immunity and increase levels of parasitemia, infection of two weeks or longer exacerbated malaria-induced morbidity and resulted in mortality in C57BL/6 mice (26). In our vaccination model, we found that chronic nematode infection of at least two weeks duration suppressed anti-OVA antibody responses to *Salmonella-*OVA (Fig. 4). Although *H. polygyrus bakeri* infection is confined to the small intestines, infection with this helminth alters gut microbial communities across the small and large intestines, and within the feces (39, 78, 79). These alterations in gut microbial communities, including enrichment in members of the order Clostridiales (39, 79) and elevated levels of their associated metabolic products, i.e. short chain fatty acids, have a significant impact on heterologous systemic immune responses (39) and likely contribute to helminth-mediated suppression of vaccine-antigen responses in our model.

We demonstrate that intestinal infection with *H. polygyrus bakeri* generated MLN-resident, polyclonal and vaccine antigen-specific, CD4^+^ T cells that produced the regulatory cytokine IL-10 (Figs. 7 and 8). Infection with *H. polygyrus bakeri* has been shown to promote the expansion of Foxp3^+^CD4^+^CD25^+^IL-10 producing regulatory T cells by producing a TGF-β mimic that can induce regulatory T cells *in vitro* even in the presence of inflammatory cytokines (80, 81). Helminth glycans have also been shown to drive regulatory T cell expansion in mixed type 2 / regulatory T cell responses characterized by the presence of IL-10-producing regulatory T cells (82). Chronic parasitic infection in mice with the systemic, blood-borne, filarial helminth *Litmosomoides sigmodontis* induces an expansion of splenic, Foxp3^-^ IL-10^+^, T regulatory 1 (Tr1) cells and reduces the quantity and quality of influenza vaccine-specific antibody responses (83). In an environmental enteric dysfunction model comprised of severely malnourished mice chronically infected with adherent *E. coli*, LP-resident Foxp3^+^RORγT^+^Tregs were associated with impaired antibody responses to an oral heat labile toxin vaccine (84). Our data demonstrate that neither a systemic chronic infection, nor severe malnutrition is required for the expansion of infection-associated regulatory T cells and suppressed vaccine-specific antibody responses. We show that in well-nourished hosts, a strictly enteric chronic helminth infection promoted the expansion of IL-10-producing T cells and impaired antibody responses to both injectable and live attenuated oral vaccines.

Our findings confirm that helminth-induced alteration of the intestinal microenvironment has systemic consequences, in this case, down-modulating immune responses to parenteral and oral vaccination. Antigen-specific antibody responses to different vaccine formulations (live attenuated *Salmonella* vaccine and protein adsorbed to alum) administered via different routes (oral and intramuscular) were suppressed by preexisting intestinal helminth infection. Our findings suggest that the immune suppressive environment generated by intestinal helminths to promote their survival impacts “third party” immune responses that may hinder the development of vaccine-induced protective immunity. Thus, the potential need to eliminate these parasites prior to vaccination should be considered when targeting populations with endemic helminth infection. In fact, large-scale clinical trials in a helminthic-endemic area (Uganda) have recently been proposed to investigate whether anti-helminthic therapies will enhance antibody and T-effector cytokine responses to both injectable and oral vaccines in school age children (61).

Future studies exploring CD4^+^T-cell independent mechanisms and their possible contributions to helminth-induced suppression of vaccine-induced antibody responses is also warranted. Our data make clear that the optimization of vaccine schedules in helminth-endemic regions must take into account that even strictly enteric helminth infection alters local mucosal and systemic immune responses to vaccination.

## Abbreviations

GALT: gut-associated lymphoid tissue
LP: lamina propria
MLN: mesenteric lymph node
OT-II Tg: OT-II transgenic
RAG1 KO: recombination-activating gene 1– deficient
Treg: regulatory T cell
Teff: T-effector cell
WHO: World Health Organization

## ACKNOWLEDGMENTS

We thank Eveline Wu, Adam Kimple, Laura Raffield, Tessa-Jonne Ropp, and Morika Williams for critical review of this manuscript. We thank Michele Ahl and the MGH Center for Comparative Medicine, Natalia Riddick and the UNC Colony Management Core in the UNC Department of Comparative Medicine, Donald W. Smith, Kabir S. Matharu, Bethany Mingle, Christian Bauer and Nikhil Milind for technical assistance.

---

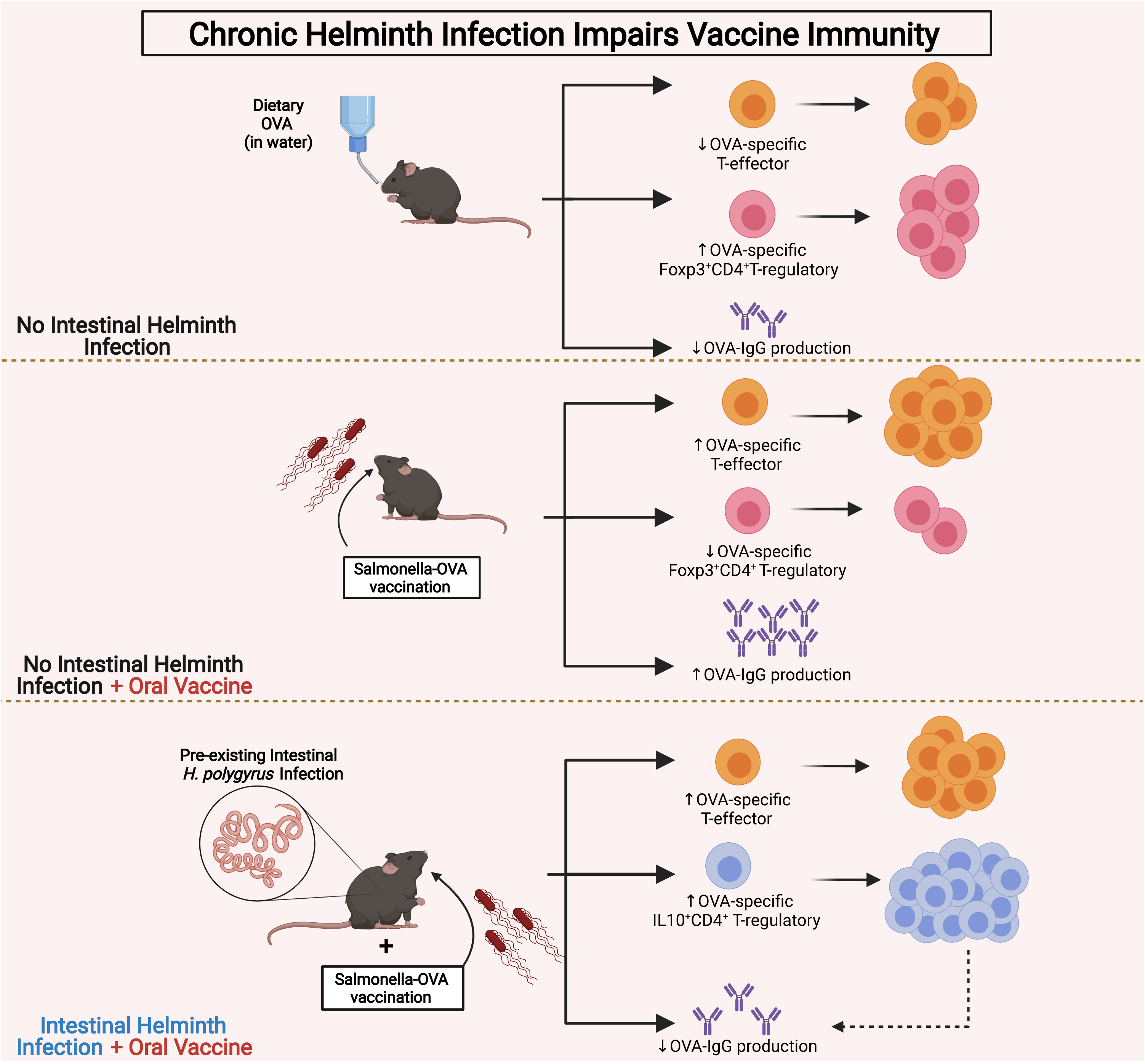

**Supplemenetary Figure 1.**
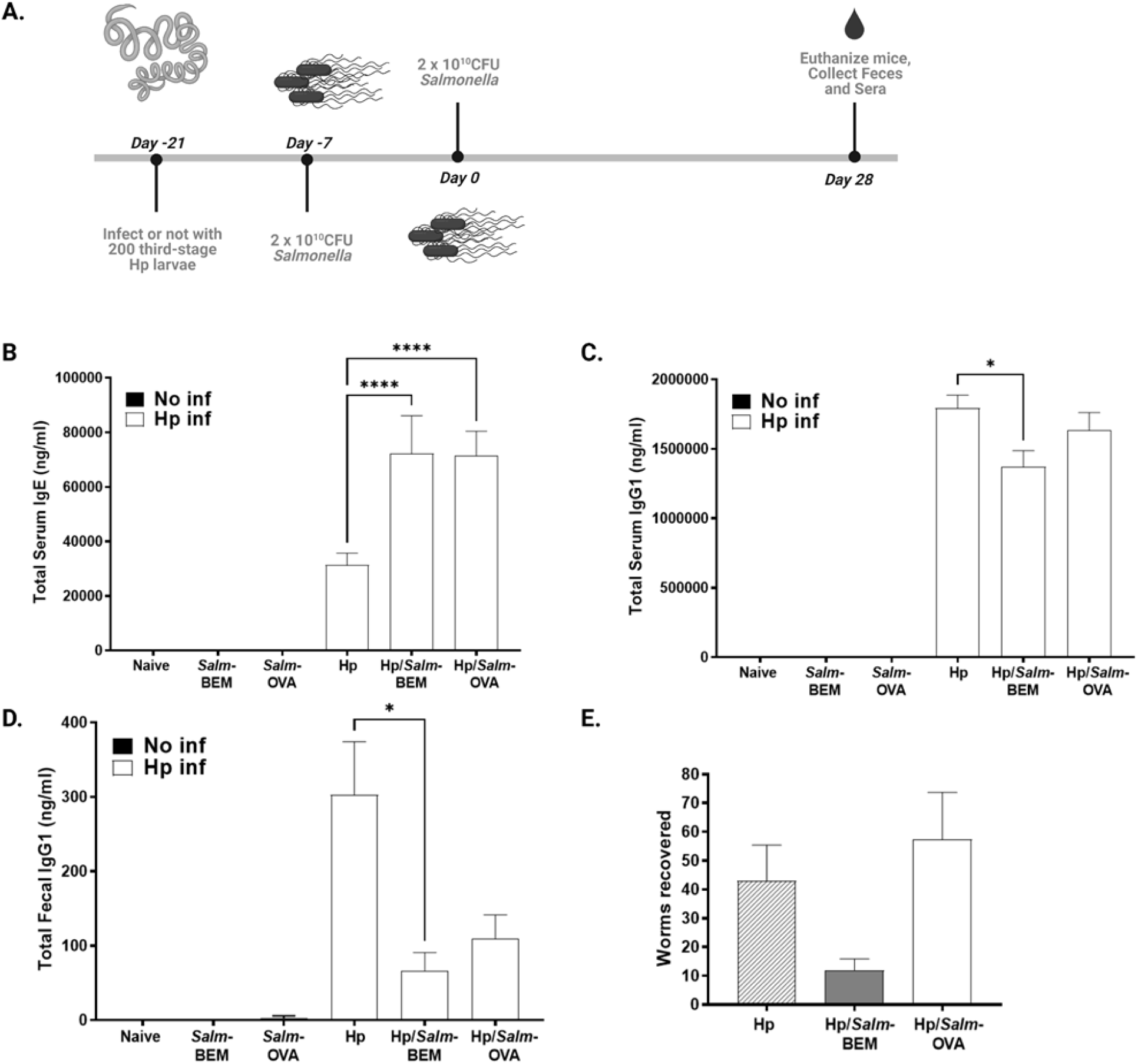

**Supplemenetary Figure 2.**
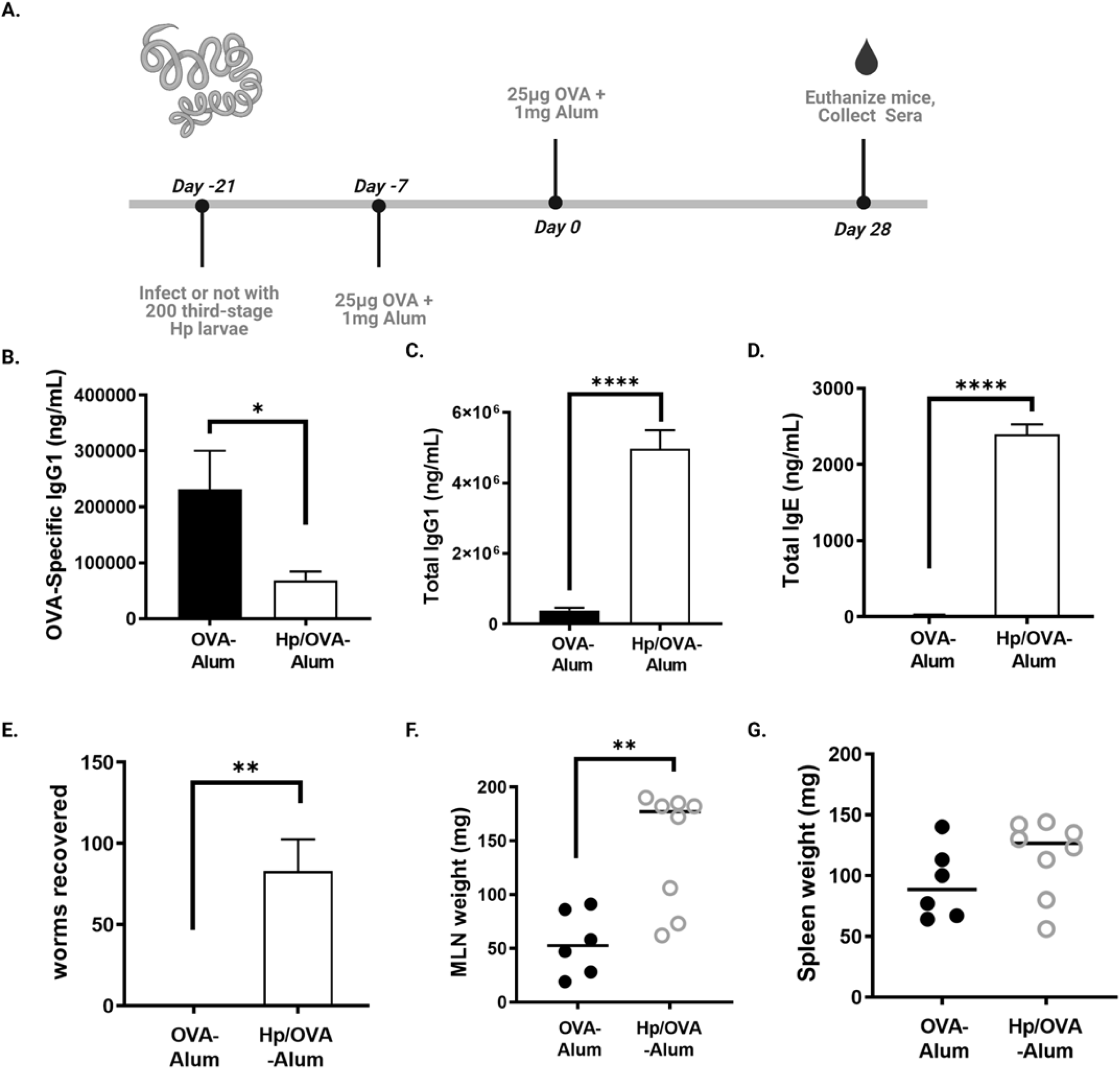

**Supplemenetary Figure 3.**
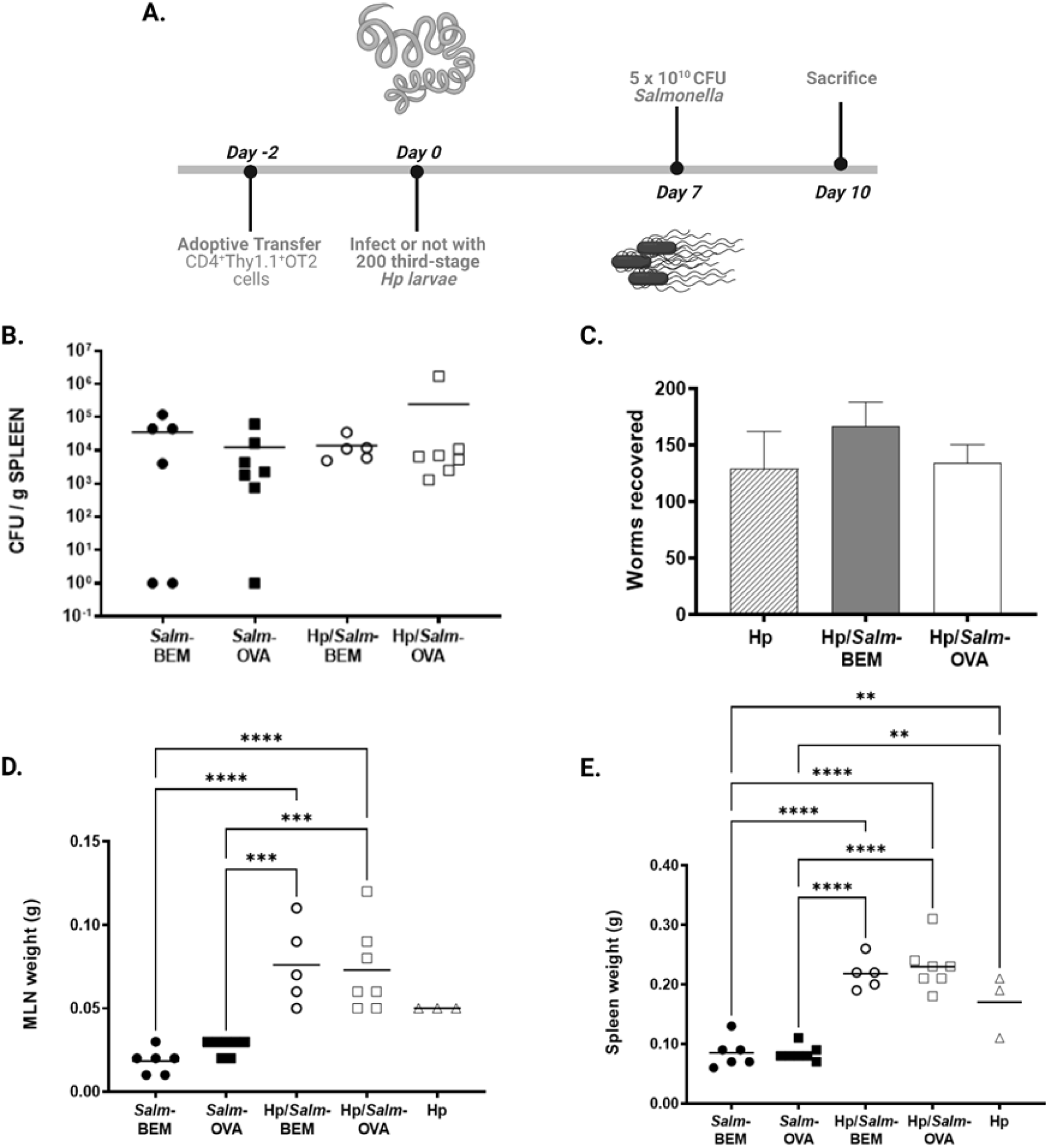

**Supplemenetary Figure 4.**
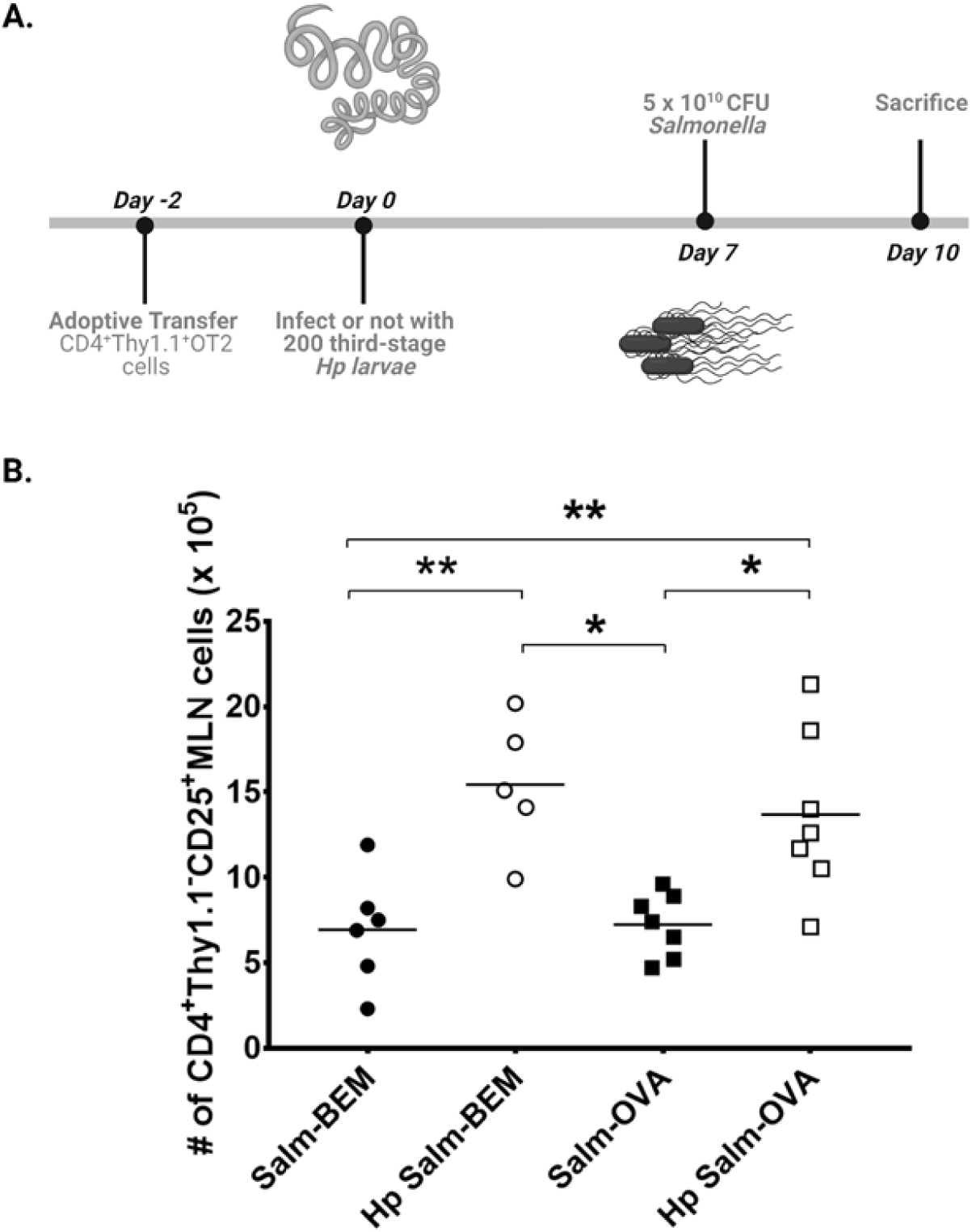

## REFERENCES

1. Centers for Disease, C., and Prevention. 2006. Vaccine preventable deaths and the Global Immunization Vision and Strategy, 2006-2015. MMWR Morb Mortal Wkly Rep 55: 511–515.

2. Pulendran, B. 2007. Tolls and beyond--many roads to vaccine immunity. N Engl J Med 356: 1776–1778.

3. Piot, P., H. J. Larson, K. L. O’Brien, J. N’Kengasong, E. Ng, S. Sow, and B. Kampmann. 2019. Immunization: vital progress, unfinished agenda. Nature 575: 119–129.

4. World Health Organization. 2020. Immunization agenda 2030: a global strategy to leave no one behind. World Health Organization, Geneva, Switzerland. p. 1–60.

5. Andre, F. E., R. Booy, H. L. Bock, J. Clemens, S. K. Datta, T. J. John, B. W. Lee, S. Lolekha, H. Peltola, T. A. Ruff, M. Santosham, and H. J. Schmitt. 2008. Vaccination greatly reduces disease, disability, death and inequity worldwide. Bull World Health Organ 86: 140–146.

6. Berkley, S. 2019. The Power of Vaccines and How Gavi Has Helped Make the World Healthier: 2019 Lasker-Bloomberg Public Service Award. Jama 322: 1251–1252.

7. Abbas, K., S. R. Procter, K. van Zandvoort, A. Clark, S. Funk, T. Mengistu, D. Hogan, E. Dansereau, M. Jit, and S. Flasche. 2020. Routine childhood immunisation during the COVID-19 pandemic in Africa: a benefit-risk analysis of health benefits versus excess risk of SARS-CoV-2 infection. Lancet Glob Health 8: e1264–e1272.

8. Curtiss, R., 3rd. 2002. Bacterial infectious disease control by vaccine development. J Clin Invest 110: 1061–1066.

9. Holmgren, J., and C. Czerkinsky. 2005. Mucosal immunity and vaccines. Nat Med 11: S45–53.

10. Pascual, D. W. 2007. Vaccines are for dinner. Proc Natl Acad Sci U S A 104: 10757–10758.

11. Redweik, G. A. J., K. Daniels, A. J. Severin, M. Lyte, and M. Mellata. 2019. Oral Treatments With Probiotics and Live Salmonella Vaccine Induce Unique Changes in Gut Neurochemicals and Microbiome in Chickens. Front Microbiol 10: 3064.

12. Redweik, G. A. J., Z. R. Stromberg, A. Van Goor, and M. Mellata. 2020. Protection against avian pathogenic Escherichia coli and Salmonella Kentucky exhibited in chickens given both probiotics and live Salmonella vaccine. Poult Sci 99: 752–762.

13. Pennington, S. H., A. L. Thompson, A. K. Wright, D. M. Ferreira, K. C. Jambo, A. D. Wright, B. Faragher, J. W. Gilmour, S. B. Gordon, and M. A. Gordon. 2016. Oral Typhoid Vaccination With Live-Attenuated Salmonella Typhi Strain Ty21a Generates Ty21a-Responsive and Heterologous Influenza Virus-Responsive CD4+ and CD8+ T Cells at the Human Intestinal Mucosa. J Infect Dis 213: 1809–1819.

14. Carreno, J. M., C. Perez-Shibayama, C. Gil-Cruz, C. Lopez-Macias, P. Vernazza, B. Ludewig, and W. C. Albrich. 2017. Evolution of Salmonella Typhi outer membrane protein-specific T and B cell responses in humans following oral Ty21a vaccination: A randomized clinical trial. PLoS One 12: e0178669.

15. Wahid, R., K. L. Kotloff, M. M. Levine, and M. B. Sztein. 2019. Cell mediated immune responses elicited in volunteers following immunization with candidate live oral Salmonella enterica serovar Paratyphi A attenuated vaccine strain CVD 1902. Clin Immunol 201: 61–69.

16. Mastroeni, P., B. Villarreal-Ramos, and C. E. Hormaeche. 1993. Adoptive transfer of immunity to oral challenge with virulent salmonellae in innately susceptible BALB/c mice requires both immune serum and T cells. Infect Immun 61: 3981–3984.

17. Hess, J., C. Ladel, D. Miko, and S. H. Kaufmann. 1996. Salmonella typhimurium aroA-infection in gene-targeted immunodeficient mice: major role of CD4+ TCR-alpha beta cells and IFN-gamma in bacterial clearance independent of intracellular location. J Immunol 156: 3321–3326.

18. Wroblewska, J. A., Y. Zhang, H. Tang, X. Guo, C. Nagler, and Y. X. Fu. 2017. Cutting Edge: Lymphotoxin Signaling Is Essential for Clearance of Salmonella from the Gut Lumen and Generation of Anti-Salmonella Protective Immunity. J Immunol 198: 55–60.

19. Mittrucker, H. W., and S. H. Kaufmann. 2000. Immune response to infection with Salmonella typhimurium in mice. J Leukoc Biol 67: 457–463.

20. Borkow, G., and Z. Bentwich. 2008. Chronic parasite infections cause immune changes that could affect successful vaccination. Trends Parasitol 24: 243–245.

21. Urban, J. F., Jr., N. R. Steenhard, G. I. Solano-Aguilar, H. D. Dawson, O. I. Iweala, C. R. Nagler, G. S. Noland, N. Kumar, R. M. Anthony, T. Shea-Donohue, J. Weinstock, and W. C. Gause. 2007. Infection with parasitic nematodes confounds vaccination efficacy. Vet Parasitol 148: 14–20.

22. Reese, T. A., K. Bi, A. Kambal, A. Filali-Mouhim, L. K. Beura, M. C. Burger, B. Pulendran, R. P. Sekaly, S. C. Jameson, D. Masopust, W. N. Haining, and H. W. Virgin. 2016. Sequential Infection with Common Pathogens Promotes Human-like Immune Gene Expression and Altered Vaccine Response. Cell Host Microbe 19: 713–719.

23. Freeman, M. C., O. Akogun, V. Belizario, Jr., S. J. Brooker, T. W. Gyorkos, R. Imtiaz, A. Krolewiecki, S. Lee, S. H. Matendechero, R. L. Pullan, and J. Utzinger. 2019. Challenges and opportunities for control and elimination of soil-transmitted helminth infection beyond 2020. PLoS Negl Trop Dis 13: e0007201.

24. Pullan, R. L., J. L. Smith, R. Jasrasaria, and S. J. Brooker. 2014. Global numbers of infection and disease burden of soil transmitted helminth infections in 2010. Parasit Vectors 7: 37.

25. Elias, D., S. Britton, A. Kassu, and H. Akuffo. 2007. Chronic helminth infections may negatively influence immunity against tuberculosis and other diseases of public health importance. Expert Rev Anti Infect Ther 5: 475–484.

26. Su, Z., M. Segura, K. Morgan, J. C. Loredo-Osti, and M. M. Stevenson. 2005. Impairment of protective immunity to blood-stage malaria by concurrent nematode infection. Infect Immun 73: 3531–3539.

27. Shi, H. N., C. J. Ingui, I. Dodge, and C. Nagler-Anderson. 1998. A helminth-induced mucosal Th2 response alters nonresponsiveness to oral administration of a soluble antigen. J. Immunol. 160: 2449–2455.

28. Shi, H. N., H. Y. Liu, and C. Nagler-Anderson. 2000. Enteric infection acts as an adjuvant for the response to a model food antigen. J. Immunol. 165: 6174–6182.

29. Bashir, M. E. H., P. Andersen, I. J. Fuss, H. N. Shi, and C. Nagler-Anderson. 2002. An enteric helminth infection protects against an allergic response to dietary antigen. J. Immunol. 169: 3284–3292.

30. Fox, J. G., P. Beck, C. A. Dangler, M. T. Whary, T. C. Wang, H. N. Shi, and C. Nagler-Anderson. 2000. Concurrent enteric helminth infection modulates inflammation and gastric immune responses and reduces helicobacter-induced gastric atrophy. Nature Medicine 6: 536–542.

31. Su, Z., M. Segura, and M. M. Stevenson. 2006. Reduced protective efficacy of a blood-stage malaria vaccine by concurrent nematode infection. Infect Immun 74: 2138–2144.

32. Da’Dara, A. A., P. J. Skelly, C. M. Walker, and D. A. Harn. 2003. A DNA-prime/protein-boost vaccination regimen enhances Th2 immune responses but not protection following Schistosoma mansoni infection. Parasite Immunol 25: 429–437.

33. Yazdanbakhsh, M., P. G. Kremsner, and R. v. Ree. 2002. Allergy, parasites and the hygiene hypothesis. Science 296: 490–494.

34. Radford-Smith, G. L. 2005. Will worms really cure Crohn’s disease? Gut 54: 6–8.

35. Maizels, R. M., and H. J. McSorley. 2016. Regulation of the host immune system by helminth parasites. J Allergy Clin Immunol 138: 666–675.

36. Yazdanbakhsh, M., A. v. d. Biggelar, and R. M. Maizels. 2001. Th2 responses without atopy: immunoregulation in chronic helminth infections and reduced allergic disease. Trends in Immunology 22: 372–377.

37. Elliott, D. E., A. Metwali, J. Leung, T. Setiawan, A. M. Blum, M. N. Ince, L. E. Bazzone, M. J. Stadecker, J. F. Urban, Jr., and J. V. Weinstock. 2008. Colonization with Heligmosomoides polygyrus suppresses mucosal IL-17 production. J Immunol 181: 2414–2419.

38. Osborne, L. C., L. A. Monticelli, T. J. Nice, T. E. Sutherland, M. C. Siracusa, M. R. Hepworth, V. T. Tomov, D. Kobuley, S. V. Tran, K. Bittinger, A. G. Bailey, A. L. Laughlin, J. L. Boucher, E. J. Wherry, F. D. Bushman, J. E. Allen, H. W. Virgin, and D. Artis. 2014. Coinfection. Virus-helminth coinfection reveals a microbiota-independent mechanism of immunomodulation. Science 345: 578–582.

39. Zaiss, M. M., A. Rapin, L. Lebon, L. K. Dubey, I. Mosconi, K. Sarter, A. Piersigilli, L. Menin, A. W. Walker, J. Rougemont, O. Paerewijck, P. Geldhof, K. D. McCoy, A. J. Macpherson, J. Croese, P. R. Giacomin, A. Loukas, T. Junt, B. J. Marsland, and N. L. Harris. 2015. The Intestinal Microbiota Contributes to the Ability of Helminths to Modulate Allergic Inflammation. Immunity 43: 998–1010.

40. Mahoney, R. T., A. Krattiger, J. D. Clemens, and R. Curtiss, 3rd. 2007. The introduction of new vaccines into developing countries. IV: Global Access Strategies. Vaccine 25: 4003–4011.

41. Iweala, O. I., D. W. Smith, K. S. Matharu, I. Sada-Ovalle, D. D. Nguyen, R. H. Dekruyff, D. T. Umetsu, S. M. Behar, and C. R. Nagler. 2009. Vaccine-induced antibody isotypes are skewed by impaired CD4 T cell and invariant NKT cell effector responses in MyD88-deficient mice. J Immunol 183: 2252–2260.

42. Bettelli, E., Y. Carrier, W. Gao, T. Korn, T. B. Strom, M. Oukka, H. L. Weiner, and V. K. Kuchroo. 2006. Reciprocal developmental pathways for the generation of pathogenic effector TH17 and regulatory T cells. Nature 441: 235–238.

43. Esterhazy, D., M. C. C. Canesso, L. Mesin, P. A. Muller, T. B. R. de Castro, A. Lockhart, M. ElJalby, A. M. C. Faria, and D. Mucida. 2019. Compartmentalized gut lymph node drainage dictates adaptive immune responses. Nature 569: 126–130.

44. Sun, C. M., J. A. Hall, R. B. Blank, N. Bouladoux, M. Oukka, J. R. Mora, and Y. Belkaid. 2007. Small intestine lamina propria dendritic cells promote de novo generation of Foxp3 T reg cells via retinoic acid. J Exp Med 204: 1775–1785.

45. Hoiseth, S. K., and B. A. Stocker. 1981. Aromatic-dependent Salmonella typhimurium are non-virulent and effective as live vaccines. Nature 291: 238–239.

46. Bryce, P. J., C. B. Mathias, K. L. Harrison, T. Watanabe, R. S. Geha, and H. C. Oettgen. 2006. The H1 histamine receptor regulates allergic lung responses. J Clin Invest 116: 1624–1632.

47. Urban, J. F., I. M. Katona, W. E. Paul, and F. D. Finkelman. 1991. Interleukin 4 is important in protective immunity to a gastrointestinal nematode infection in mice. *Proc. Natl. Acad. Sci.*, USA 83: 5513–5517.

48. Setiawan, T., A. Metwali, A. M. Blum, M. N. Ince, J. F. Urban, Jr., D. E. Elliott, and J. V. Weinstock. 2007. Heligmosomoides polygyrus promotes regulatory T-cell cytokine production in the murine normal distal intestine. Infect Immun 75: 4655–4663.

49. Kaji, H., M. Kawada, A. Tai, H. Kanzaki, and I. Yamamoto. 2003. Augmentation by Bursaphelenchus xylophilus, a pine wood nematode, of polyclonal IgE production induced by lipopolysaccharide plus interleukin-4 in murine splenocytes. J Pharmacol Sci 91: 158–162.

50. Cerutti, A. 2008. The regulation of IgA class switching. Nat Rev Immunol 8: 421–434.

51. Shevach, E. M. 2002. CD4+CD25+ suppressor T cells: more questions than answers. Nat. Rev. Immunol. 2: 389–400.

52. Finney, C. A., M. D. Taylor, M. S. Wilson, and R. M. Maizels. 2007. Expansion and activation of CD4(+)CD25(+) regulatory T cells in Heligmosomoides polygyrus infection. Eur J Immunol 37: 1874–1886.

53. Hang, L., S. Kumar, A. M. Blum, J. F. Urban, Jr., M. C. Fantini, and J. V. Weinstock. 2019. Heligmosomoides polygyrus bakeri Infection Decreases Smad7 Expression in Intestinal CD4(+) T Cells, Which Allows TGF-beta to Induce IL-10-Producing Regulatory T Cells That Block Colitis. J Immunol 202: 2473–2481.

54. Mangan, N. E., N. van Rooijen, A. N. McKenzie, and P. G. Fallon. 2006. Helminth-modified pulmonary immune response protects mice from allergen-induced airway hyperresponsiveness. J Immunol 176: 138–147.

55. Su, H., Q. Liu, X. Bian, S. Wang, R. Curtiss, 3rd, and Q. Kong. 2021. Synthesis and delivery of Streptococcus pneumoniae capsular polysaccharides by recombinant attenuated Salmonella vaccines. Proc Natl Acad Sci U S A 118.

56. Lin, Z., H. Sun, Y. Ma, X. Zhou, H. Jiang, X. Wang, J. Song, Z. Tang, Q. Bian, Z. Zhang, Y. Huang, and X. Yu. 2020. Evaluation of immune response to Bacillus subtilis spores expressing Clonorchis sinensis serpin3. Parasitology 147: 1080–1087.

57. Wang, L., X. Wang, K. Bi, X. Sun, J. Yang, Y. Gu, J. Huang, B. Zhan, and X. Zhu. 2016. Oral Vaccination with Attenuated Salmonella typhimurium-Delivered TsPmy DNA Vaccine Elicits Protective Immunity against Trichinella spiralis in BALB/c Mice. PLoS Negl Trop Dis 10: e0004952.

58. Srinivasa Reddy, Y. K. Narendra Babu, P. Uday Kumar, N. Harishankar, S. Qadri, M. V. Surekha, R. Hemalatha, and B. Dinesh Kumar. 2021. Nonclinical safety evaluation of oral recombinant anti-human papilloma virus vaccine (RHPV 16 & 18): Regulatory toxicology studies in mice, rats and rabbits - An innovative approach. Vaccine 39: 853–863.

59. Liu, Q., H. Su, X. Bian, S. Wang, and Q. Kong. 2020. Live attenuated Salmonella Typhimurium with monophosphoryl lipid A retains ability to induce T-cell and humoral immune responses against heterologous polysaccharide of Shigella flexneri 2a. Int J Med Microbiol 310: 151427.

60. Kabagenyi, J., A. Natukunda, J. Nassuuna, R. E. Sanya, M. Nampijja, E. L. Webb, A. M. Elliott, and G. Nkurunungi. 2020. Urban-rural differences in immune responses to mycobacterial and tetanus vaccine antigens in a tropical setting: A role for helminths? Parasitol Int 78: 102132.

61. Nkurunungi, G., L. Zirimenya, J. Nassuuna, A. Natukunda, P. N. Kabuubi, E. Niwagaba, G. Oduru, G. Kabami, R. Amongin, A. Mutebe, M. Namutebi, C. Zziwa, S. Amongi, C. Ninsiima, C. Onen, F. Akello, M. Sewankambo, S. Kiwanuka, R. Kizindo, J. Kaweesa, S. Cose, E. Webb, A. M. Elliott, P. t. team, and P. t. t. p. investigator. 2021. Effect of intensive treatment for schistosomiasis on immune responses to vaccines among rural Ugandan island adolescents: randomised controlled trial protocol A for the ‘POPulation differences in VACcine responses’ (POPVAC) programme. BMJ Open 11: e040426.

62. Clark, C. E., M. P. Fay, M. E. Chico, C. A. Sandoval, M. G. Vaca, A. Boyd, P. J. Cooper, and T. B. Nutman. 2016. Maternal Helminth Infection Is Associated With Higher Infant Immunoglobulin A Titers to Antigen in Orally Administered Vaccines. J Infect Dis 213: 1996–2004.

63. Hallander, H. O., M. Paniagua, F. Espinoza, P. Askelof, E. Corrales, M. Ringman, and J. Storsaeter. 2002. Calibrated serological techniques demonstrate significant different serum response rates to an oral killed cholera vaccine between Swedish and Nicaraguan children. Vaccine 21: 138–145.

64. Jiang, V., B. Jiang, J. Tate, U. D. Parashar, and M. M. Patel. 2010. Performance of rotavirus vaccines in developed and developing countries. Hum Vaccin 6: 532–542.

65. Patriarca, P. A., P. F. Wright, and T. J. John. 1991. Factors affecting the immunogenicity of oral poliovirus vaccine in developing countries: review. Rev Infect Dis 13: 926–939.

66. Sabin, E. A., M. I. Araujo, E. M. Carvalho, and E. J. Pearce. 1996. Impairment of tetanus toxoid-specific Th1-like immune responses in humans infected with Schistosoma mansoni. J Infect Dis 173: 269–272.

67. Cooper, P. J., M. Chico, C. Sandoval, I. Espinel, A. Guevara, M. M. Levine, G. E. Griffin, and T. B. Nutman. 2001. Human infection with Ascaris lumbricoides is associated with suppression of the interleukin-2 response to recombinant cholera toxin B subunit following vaccination with the live oral cholera vaccine CVD 103-HgR. Infect Immun 69: 1574–1580.

68. Elias, D., S. Britton, A. Aseffa, H. Engers, and H. Akuffo. 2008. Poor immunogenicity of BCG in helminth infected population is associated with increased in vitro TGF-beta production. Vaccine 26: 3897–3902.

69. Elias, D., D. Wolday, H. Akuffo, B. Petros, U. Bronner, and S. Britton. 2001. Effect of deworming on human T cell responses to mycobacterial antigens in helminth-exposed individuals before and after bacille Calmette-Guerin (BCG) vaccination. Clin Exp Immunol 123: 219–225.

70. Hunter, M. M., A. Wang, C. L. Hirota, and D. M. McKay. 2005. Neutralizing anti-IL-10 antibody blocks the protective effect of tapeworm infection in a murine model of chemically induced colitis. J Immunol 174: 7368–7375.

71. McConchie, B. W., H. H. Norris, V. G. Bundoc, S. Trivedi, A. Boesen, J. F. Urban, Jr., and A. M. Keane-Myers. 2006. Ascaris suum-derived products suppress mucosal allergic inflammation in an interleukin-10-independent manner via interference with dendritic cell function. Infect Immun 74: 6632–6641.

72. Wilson, M. S., M. D. Taylor, A. Balic, C. A. Finney, J. R. Lamb, and R. M. Maizels. 2005. Suppression of allergic airway inflammation by helminth-induced regulatory T cells. J Exp Med 202: 1199–1212.

73. Nookala, S., S. Srinivasan, P. Kaliraj, R. B. Narayanan, and T. B. Nutman. 2004. Impairment of tetanus-specific cellular and humoral responses following tetanus vaccination in human lymphatic filariasis. Infect Immun 72: 2598–2604.

74. Jordan, M. B., D. M. Mills, J. Kappler, P. Marrack, and J. C. Cambier. 2004. Promotion of B cell immune responses via an alum-induced myeloid cell population. Science 304: 1808–1810.

75. Serre, K., E. Mohr, K. M. Toellner, A. F. Cunningham, S. Granjeaud, R. Bird, and I. C. MacLennan. 2008. Molecular differences between the divergent responses of ovalbumin-specific CD4 T cells to alum-precipitated ovalbumin compared to ovalbumin expressed by Salmonella. Mol Immunol 45: 3558–3566.

76. Kilian, H. D., and G. Nielsen. 1989. Cell-mediated and humoral immune response to tetanus vaccinations in onchocerciasis patients. Trop Med Parasitol 40: 285–291.

77. Lynch, N. R., R. I. Lopez, M. C. Di Prisco-Fuenmayor, I. Hagel, L. Medouze, G. Viana, C. Ortega, and G. Prato. 1987. Allergic reactivity and socio-economic level in a tropical environment. Clin Allergy 17: 199–207.

78. Rapin, A., and N. L. Harris. 2018. Helminth-Bacterial Interactions: Cause and Consequence. Trends Immunol 39: 724–733.

79. Ramanan, D., R. Bowcutt, S. C. Lee, M. S. Tang, Z. D. Kurtz, Y. Ding, K. Honda, W. C. Gause, M. J. Blaser, R. A. Bonneau, Y. A. Lim, P. Loke, and K. Cadwell. 2016. Helminth infection promotes colonization resistance via type 2 immunity. Science 352: 608–612.

80. Rausch, S., J. Huehn, D. Kirchhoff, J. Rzepecka, C. Schnoeller, S. Pillai, C. Loddenkemper, A. Scheffold, A. Hamann, R. Lucius, and S. Hartmann. 2008. Functional analysis of effector and regulatory T cells in a parasitic nematode infection. Infect Immun 76: 1908–1919.

81. Grainger, J. R., K. A. Smith, J. P. Hewitson, H. J. McSorley, Y. Harcus, K. J. Filbey, C. A. Finney, E. J. Greenwood, D. P. Knox, M. S. Wilson, Y. Belkaid, A. Y. Rudensky, and R. M. Maizels. 2010. Helminth secretions induce de novo T cell Foxp3 expression and regulatory function through the TGF-beta pathway. J Exp Med 207: 2331–2341.

82. Rodriguez, E., P. Carasi, S. Frigerio, V. da Costa, S. van Vliet, V. Noya, N. Brossard, Y. van Kooyk, J. J. Garcia-Vallejo, and T. Freire. 2017. Fasciola hepatica Immune Regulates CD11c(+) Cells by Interacting with the Macrophage Gal/GalNAc Lectin. Front Immunol 8: 264.

83. Hartmann, W., M. L. Brunn, N. Stetter, N. Gagliani, F. Muscate, S. Stanelle-Bertram, G. Gabriel, and M. Breloer. 2019. Helminth Infections Suppress the Efficacy of Vaccination against Seasonal Influenza. Cell Rep 29: 2243–2256 e2244.

84. Bhattacharjee, A., A. H. P. Burr, A. E. Overacre-Delgoffe, J. T. Tometich, D. Yang, B. R. Huckestein, J. L. Linehan, S. P. Spencer, J. A. Hall, O. J. Harrison, D. Morais da Fonseca, E. B. Norton, Y. Belkaid, and T. W. Hand. 2021. Environmental enteric dysfunction induces regulatory T cells that inhibit local CD4+ T cell responses and impair oral vaccine efficacy. Immunity 54: 1745–1757 e1747.

